# Ethanolamine-phosphate on the second mannose is the preferential bridge for some of the brain GPI-anchored proteins

**DOI:** 10.1101/2020.11.26.399477

**Authors:** Mizuki Ishida, Yuta Maki, Akinori Ninomiya, Yoko Takada, Philippe Campeau, Taroh Kinoshita, Yoshiko Murakami

## Abstract

Glycosylphosphatidylinositols (GPIs) are glycolipids that anchor many proteins (GPI-APs) on the cell surface. The core glycan of GPI precursor has three mannoses, which in mammals, are all modified by ethanolamine-phosphate (EthN-P). It is postulated that EthN-P on the third mannose (EthN-P-Man3) is the bridge between GPI to the protein and the second (EthN-P-Man2) is removed after GPI-protein attachment. However, EthN-P-Man2 may not be always transient, as mutations of PIGG, the enzyme that transfers EthN-P to Man2, result in inherited GPI deficiencies (IGDs), characterized by neuronal dysfunctions. Here, we show EthN-P on Man2 is the preferential bridge in some GPI-APs, among them, the ect-5’-nucleotidase and netrin G2. We found that CD59, a GPI-AP, is attached via EthN-P-Man2 both in *PIGB*-knockout cells, in which GPI lacks Man3 and with a small fraction, in wild type cells. Our findings modify the current view of GPI anchoring and provide mechanistic bases of IGDs caused by *PIGG* mutations.

## Introduction

Glycosylphosphatidylinositols (GPIs) are glycolipids found ubiquitously as membrane anchors of many proteins in eukaryotic organisms. The structures of GPI anchors were first determined in rat Thy-1 and trypanosome variant surface glycoprotein in the 1980s (Homans et al., 1988; Masterson et al., 1989), and shown that its backbone structure is highly conserved among all eukaryotes (Ferguson et al., 2015). Its basic structure is

EthN-P-6-Man-α1,2-Man-α1,6-Man-α1,4-GlcN-α1,6-*myo*Inositol-phospholipid, where GlcN, Man, and EthN-P are glucosamine, mannose, and ethanolamine-phosphate respectively (Homans et al., 1988). This backbone is modified by various side chain groups in different organisms, or even in different proteins of the same organism. An example of the latter is mammalian GPI-APs, in which the first α1,4-linked Man (Man1) is always modified by EthN-P side chain attached on the 2-position, but a fourth Man may be attached to the third Man (Man3) via α1,2-linkage.

GPI anchor precursors and precursor proteins are synthesized separately, and assembly of GPI-anchored proteins (GPI-APs) occurs in the endoplasmic reticulum (ER). The GPI transamidase complex recognizes the C-terminal GPI attachment signal of precursor proteins (Muniz and Riezman, 2016), processes and attaches them to EthN-P on Man3 of GPI via an amide bond (Benghezal et al., 1996; Hamburger et al., 1995). This EthN-P is thus called “bridging EthN-P”. Both glycan and lipid moieties of GPI-APs are further modified in the ER and the Golgi before selectively trafficked to the cell surface. In mammalian cells, at least 27 genes, termed *PIG* (Phosphatidylinositolglycan) and *PGAP* (Post GPI attachment to proteins) genes, are involved in the biosynthesis and maturation of GPI-APs (Kinoshita, 2020).

In humans, more than 150 GPI-APs have been characterized, which functions include enzymes, receptors, adhesion molecules and complement regulatory proteins (UniProt, 2015). Many of these GPI-APs and enzymes involved in their synthesis, play important roles in neuronal tissues. For instance, defects in *PIG* and *PGAP* genes cause inherited GPI deficiency (IGD), the main symptoms of which are intellectual disability, developmental delay, and seizures (Kuki, 2013). At present, IGD caused by mutations in 22 genes involved in GPI biosynthesis (Almeida et al., 2006; Chiyonobu et al., 2014; Edvardson et al., 2017; Ilkovski et al., 2015; Johnston et al., 2012; Johnstone et al., 2017; Knaus et al., 2019; Krawitz et al., 2012; Krawitz et al., 2010; Kvarnung et al., 2013; Makrythanasis et al., 2016; Martin et al., 2014; Maydan et al., 2011; Murakami et al., 2019; Ng et al., 2012; Nguyen et al., 2020; Nguyen et al., 2017; Nguyen et al., 2018; Pagnamenta et al., 2018) and the GPI-AP maturation pathway (Howard MF, 2014; Krawitz et al., 2013

; Murakami et al., 2014) have been reported. Defects in these genes lead mostly to embryonic death (Nozaki et al., 1999) but the vast majority of survivors, who show symptoms of IGD, have partial deficiencies in GPI biosynthesis. PIGO is an EthN-P transferase involved in attachment of the bridging EthN-P to Man3 (Hong et al., 2000), which is transferred by the mannosyltransferase PIGB to the GPI precursor (Takahashi et al., 1996). When analyzing HEK293 cells with the *PIGO* or *PIGB* gene knocked out (*PIGO*-KO or *PIGB*-KO), we unexpectedly found some GPI-APs on the surface of the cells, such as CD59 and DAF, which in turn were completely obliterated by further knockout of *PIGG*. PIGG is another EthN-P transferase involved in the attachment of the EthN-P to the second Man (Man2) in the precursor GPI (Shishioh et al., 2005), believed to be removed soon after GPI is attached to the precursor protein (Fujita et al., 2009). The physiological significance of this transient modification has been unclear (Shishioh et al., 2005). Interesting though, in *PIGG*-KO cells, CD59 and DAF, are normally transported and expressed with the canonical structure. Notwithstanding this fact, *PIGG* mutations in humans result in the mentioned IGD symptoms, suggesting an important role of PIGG in neuronal physiology (Makrythanasis et al., 2016; Tremblay-Laganière et al., 2021; Zhao et al., 2017).

In this work, we analyzed by mass spectrometry, the GPI structure of CD59 expressed in *PIGB*-KO cells and found that the EthN-P on Man2 is the bridging EthN-P for this protein. We also found CD59 and the ect-5’-nucleotidase with this same GPI structure in wild type HEK293 cells. It has long been postulated that in mammalian cells, the GPI-APs are always linked through the EthN-P on Man3. Our current data challenges this view.

## Results

### PIGG-dependent, low level expression of GPI-anchored CD59 and DAF in PIGO-KO and PIGB-KO cells

It has been postulated that PIGG-dependent, Man2 linked EthN-P is a transient structure that is removed soon after attachment of protein to the Man3 linked EthN-P. In support of this notion is the similar expression levels of the GPI-APs, CD59 and DAF, observed in *PIGG*-KO HEK293 to those in non-mutated HEK293 (wild type) cells, as determined by fluorescence-activated cell sorting (FACS) analysis (Figure 1A). PIGO dependent Man3-linked EthN-P has been thought to act as the bridge in all GPI-APs (Hong et al., 2000). Unexpectedly, however, *PIGO*-KO HEK293 cells expressed, albeit low, clearly detectable levels of CD59 and DAF, being 6% and 19% of the levels observed in wild type HEK293 cells, respectively (Figure 1B). Furthermore, the residual expression of CD59 as well as of DAF, completely disappears after further knockout of *PIGG* in *PIGO*-KO (*PIGO/PIGG*-DKO) cells, suggesting an important role of PIGG in the expression of these residual GPI-APs (Figure 1C). PIGO and PIGG are EthN-P transferases that add EthN-P to the 6-position of Man3 and Man2, respectively (Hong et al., 2000; Shishioh et al., 2005). To test the possibility that PIGG compensated the PIGO loss and added the EthN-P to Man3 in *PIGO*-KO cells, PIGG was overexpressed in the *PIGO/PIGG*-DKO cells by transfection of *PIGG* cDNA (rescuer *PIGG*; Table S3). CD59 and DAF expression was restored up to, but not above, their levels in *PIGO*-KO cells (Figure 1C), suggesting a mechanism other than compensation by PIGG. Furthermore, in *PIGO*-KO cells, CD59 was slightly sensitive, and DAF completely resistant, to bacterial phosphatidylinositol-specific phospholipase C (PI-PLC) treatment, whereas in wild type cells both proteins were highly sensitive (Figure 1D), suggesting different structures in the GPI-anchors of these two GPI-APs in *PIGO*-KO vs wild type cells (see Discussion).

**Figure 1.**
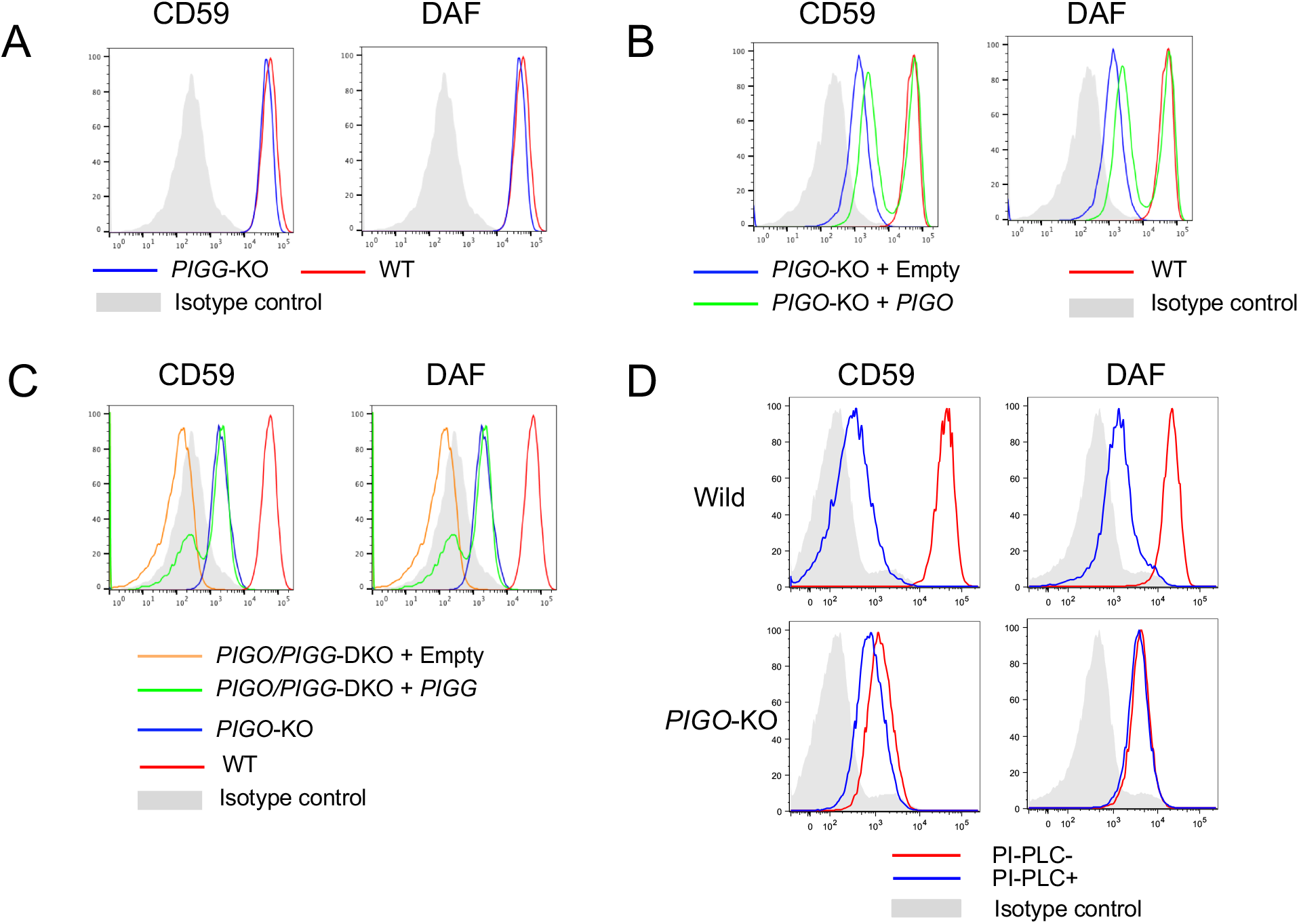
Complete silencing of residual expression of CD59 and DAF GPI-APs in *PIGO*-KO HEK293 cells by knocking out *PIGG* (*PIGO*-*PIGG*-DKO). A. Regular levels of CD59 and DAF in *PIGG*-KO cells as determined by FACS analysis. B. *PIGO*-KO cells expressing low levels of CD59 and DAF and restored by transfection of *PIGO* cDNA. C. Complete removal of residual expression of CD59 and DAF in *PIGO*-KO cells by further knockout of *PIGG* (*PIGO-PIGG*-DKO). D. Resistance of CD59 and DAF expressed in *PIGO*-KO cells to PI-PLC treatment. Isotype control, isotypic cells stained with non-matching antibody, as negative control.

Furthermore, since Man3 is transferred by PIGB to Man2 in the α1, 2 linkage way (Oriol et al., 2002; Takahashi et al., 1996), such transfer should not occur in *PIGB*-KO cells. Nevertheless, we observed low levels of CD59 and DAF in these cells, their levels being 3% and 7% to those in wild type cells, respectively (Figure 2A). In some cases, PIGZ transfers a fourth mannose (Man4) to Man3 in the same α1,2 linkage (Taron et al., 2004). Since *PIGB*-KO HEK293 cells express PIGZ, a possible explanation for the low expression of CD59 and DAF observed in these cells is that PIGZ is compensating the missing PIGB function. However, further *PIGZ* knockout in *PIGB*-KO cells did not obliterate the low levels of CD59 and DAF (Figure 2B). And again, *PIGG* knockout in *PIGB*-KO cells resulted in the disappearance of the residual CD59 and DAF, further suggesting an important role of PIGG in the residual expression of these proteins (Figure 2C). In addition, differently to those in *PIGO*-KO cells, CD59 and DAF residually expressed in *PIGB*-KO cells were more sensitive to PI-PLC, especially CD59, of which about 90% became detached from *PIGB*-KO cells surface upon PI-PLC treatment (Figure 2D). The results described above strongly suggest that the EthN-P on Man2 is also used, albeit at low levels, as bridge in the anchoring of CD59 and DAF.

**Figure 2.**
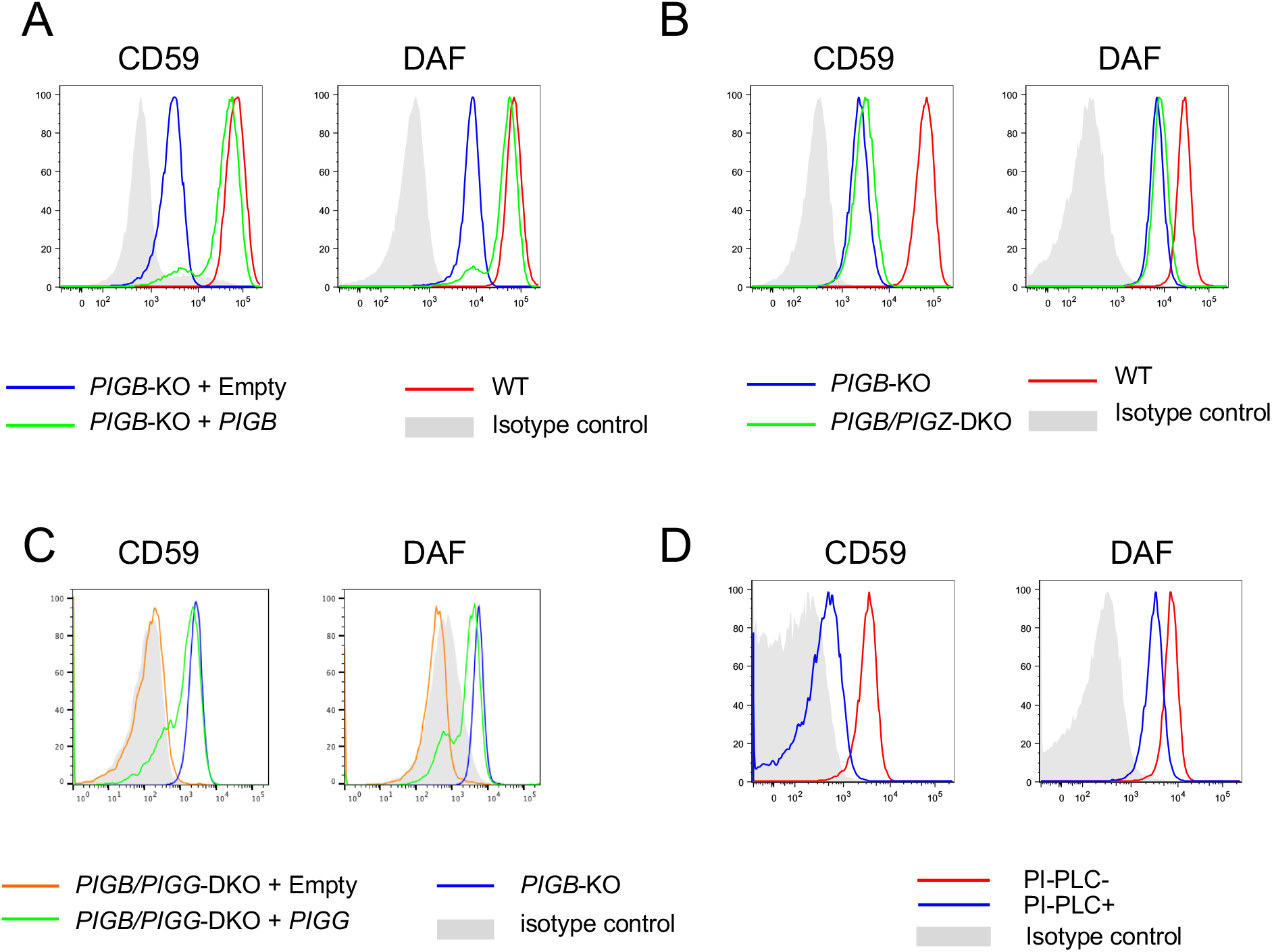
Complete silencing of residual expression of synthesized in *PIGB*-KO HEK293 cells by knocking out *PIGG* (*PIGB*-*PIGG*-DKO). A. *PIGB*-KO cells expressing low levels of CD59 and DAF, and restored by transfection of *PIGB* cDNA. B. Further knockout of *PIGZ* of *PIGB*-KO (*PIGB-PIGZ*-DKO) cells resulting in no decrease of CD59 and DAF expression levels. C. Further knockout of *PIGG* of PIGB-KO (*PIGB-PIGG*-DKO) cells resulting in complete removal of residual expression of CD59 and DAF. D. Sensitivity to PI-PLC treatment of CD59 and DAF expressed in *PIGB*-KO cells.

### Mass spectrometry (MS) reveals attachment of CD59 to Man2-linked EthN-P in PIGB-KO cells

To determine the structural basis of PIGG-dependent CD59, we stably transfected HEK293 cells with the pME plasmid vector (Kitamura et al., 1991) expressing HFGF-CD59 (pME-puro-*HFGF-CD59*, the *CD59* cDNA fused to epitope tags and GST to aid in its purification, Table S3) and highly expressing cells were isolated by cell sorting. We then knocked out *PIGB* in these cells and isolated various cell clones. HFGF-CD59 proteins were solubilized by PI-PLC-treatment from both the original HEK293-*HFGF-CD59* cells and its *PIGB*-KO cell clone, followed by GST affinity purification, SDS-PAGE and in-gel trypsin digestion (Figure S1). The C-terminal peptide of HEK293-HFGF-CD59 linked to GPI glycan from both HEK293-*HFGF-CD59* cells and their *PIGB*-KO cells, was then structurally analyzed by LC-ESI-MS.

In the HEK293-*HFGF-CD59* cells sampling, we found precursor ions corresponding to the C-terminal 11 amino acid peptide (DLCNFNEQLEN) linked to GPI with three Man and a HexNAc side chain (*m/z* 912.0^3+^ and 1367.97^2+^, MW 2733.94) (Figures 3A and S2). Analyses by MS/MS showed a group of fragment ions diagnostic of GPI (Figures 3B; *m/z* 422^+^, GlcN-Ino-P; 447^+^, EthN-P-Man-GlcN; 707^+^, EthN-P-Man-GlcN-Ino-P; 812^+^, Man-(EthN-P-)Man-GlcN-HexNAc; 851^+^, P-Man-Man-(EthN-P-)Man-GlcN; 869^+^, Man-(EthN-P-)Man-GlcN-Ino-P; 1072+, Man-(EthN-P-)Man-GlcN-Ino-P-HexNAc; 1111^+^, P-Man-Man-(EthN-P-)Man-GlcN-Ino-P; 1314^+^, P-Man-Man-(EthN-P-)Man-GlcN-Ino-P-Hex; 1421^+^, peptide-EthN; 1663^+^, peptide-EthN-P-Man; 1825^+^, peptide-EthN-P-Man-Man; 1987^+^, peptide-EthN-P-Man-Man-Man) clearly defining the structure, DLCNFNEQLEN-EthN-P-Man-Man-(EthN-P-)(HexNAc-)Man-GlcN-Ino-P- (Figure 3A). HexNAc, N-acetylhexosamine, corresponds to GalNAc in mammalian GPIs, as described (Hirata et al., 2018). We also found precursor ions of *m/z* 966.0^3+^ and 1448.5^2+^ (MW 2895.0) corresponding to a GPI-peptide having one extra Hex. The Hex could correspond to either Man4 or Gal extension of the GalNAc side chain (Figure S2) (Wang et al., 2020). No other precursor ion corresponding to the C-terminal peptide, and having any other known GPI structure, such as one lacking the GalNAc side chain, was detected. Quantitative analysis of peak area of total ion chromatogram (TIC) of each fragment, showed that most, if not all, GPIs of HFGF-CD59 expressed in wild type HEK293 cells have a GalNAc side chain and that nearly 40% of them had either Man4 or Gal extension (Figure S2).

**Figure 3.**
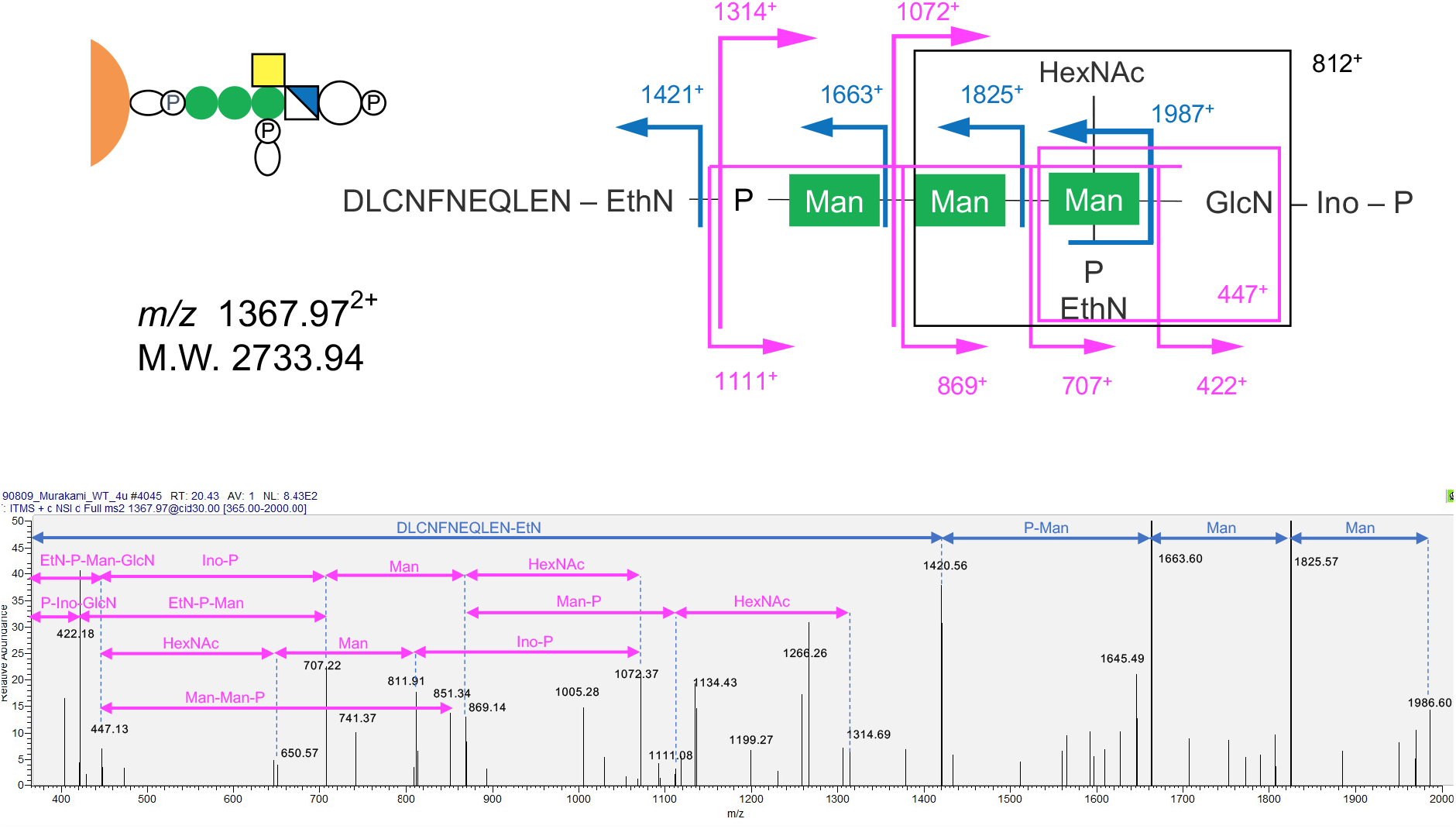
Mass spectrometric analysis of trypsin fragments of CD59 expressed in wild type HEK293 cells. A. Schematic MS/MS profile showing the CD59 C-terminal eleven amino acids linked to the typical GPI glycan, EthN-P-Man-Man- (EthN-P-)(HexNAc-)Man-GlcN-Ino-P. M.W. was calculated from the value of *m/z*. B. The MS/MS profile of a major precursor ion of *m/z* 1367.97^2+^ corresponding to the glycan shown in A.

In the *PIGB*-KO HEK293-*HFGF-CD59* cells sampling, we found precursor ions corresponding to the C-terminal 11 amino acid peptide linked to GPIs having two Man and a HexNAc side chain (*m/z* 858.0^3+^ and 1286.94^2+^; MW 2571.88) as well as some bearing Hex extension (*m/z* 912.0^3+^; MW 2733.0) (Figures 4A and S3). MS/MS analysis indicated not only the GPI diagnostic fragments (*m/z* 422^+^, 447^+^, 707^+^, 1421^+^ and 1663^+^) but also four fragments corresponding to P-Man-(EthN-P-)Man-GlcN (*m/z* 689^+^), P-Man-(EthN-P-)Man-GlcN-Ino-P (*m/z* 949^+^), P-Man-(EthN-P-)Man-GlcN-Ino-P-HexNAc (*m/z* 1152^+^), peptide-EthN-P-Man-(EthN-P-)Man (*m/z* 1948^+^) (Figure 4B). The P-Man-(EthN-P-)Man structure can only be derived from a GPI core with the protein attached to the EthN-P on Man2. Results of TIC analysis showed that most GPIs of HFGF-CD59 in *PIGB*-KO HEK293 cells have only two Man and the GalNAc side chain, with approximately 20% bearing the Gal extension (Figure S3).

**Figure 4.**
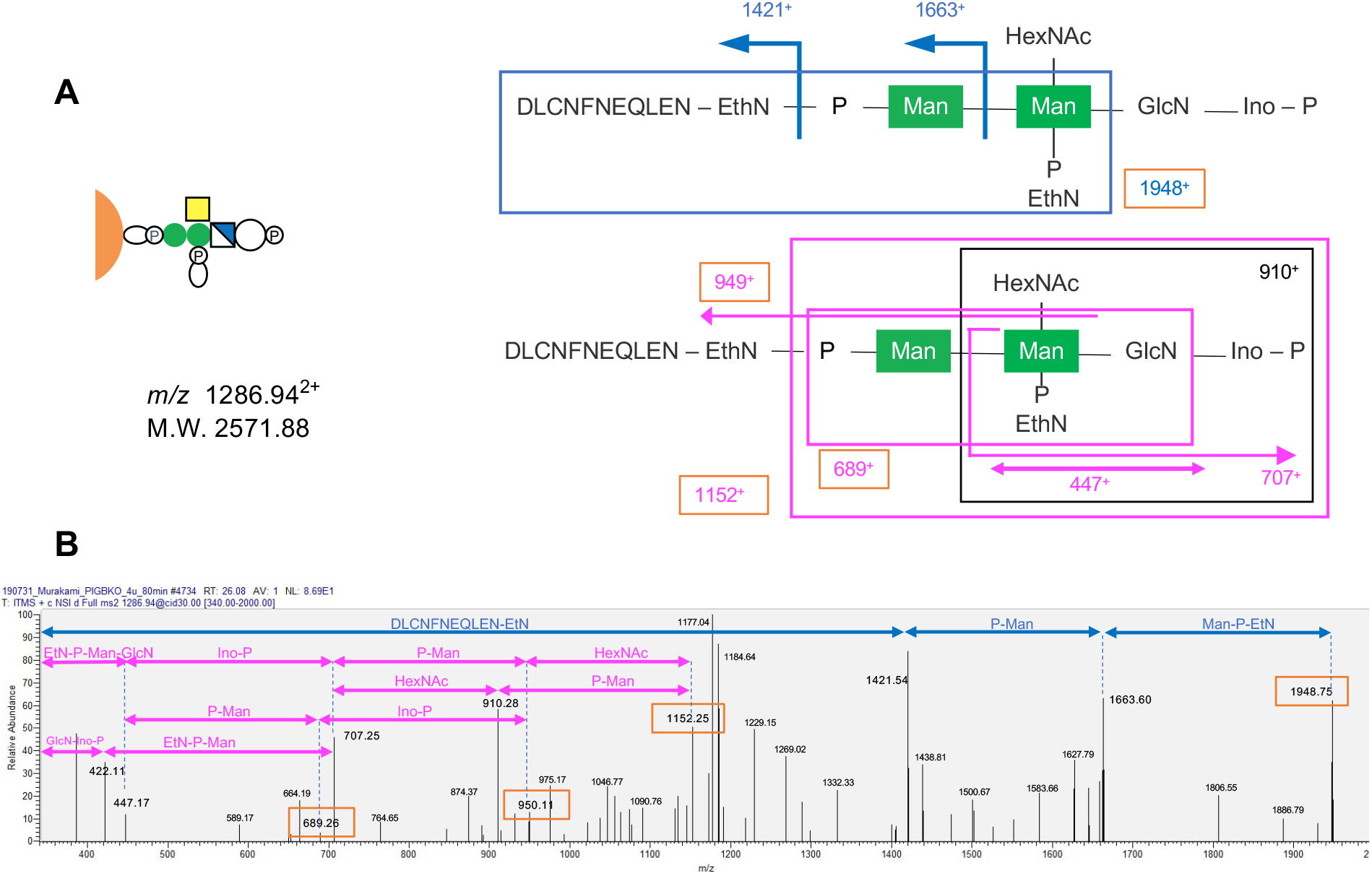
Mass spectrometric analysis of trypsin fragments of CD59 expressed in *PIGB*-KO HEK293 cells. A, Schematic MS/MS profile showing the CD59 C-terminal eleven amino acids linked to a GPI glycan with only two mannoses. M.W. was calculated from the value of *m/z*. B. MS/MS analysis of a precursor ion of *m/z* 1286.94^2+^. Fragment ions *m/z* 689^+^ corresponding to -P-Man-(EthN-P-)Man-GlcN, m/z 949^+^ corresponding to P-Man-(EthN-P-)Man-GlcN-Ino-P, *m/z* 1152^+^ corresponding to P-Man-(EthN-P-)Man-GlcN-Ino-P-HexNAc and *m/z* 1948^+^ corresponding to (11 amino acids)-EthN-P-Man-(EthN-P-)Man, indicate CD59 C-terminus attachment to EthN-P linked to the second mannose.

### *A small fraction of HFGF-CD59 in HEK293-*HFGF-CD59 *cells is attached to Man2-linked EthN-P in GPI*

We next determined whether CD59 protein attachment to Man2-linked EthN-P occurs in wild type cells by reanalyzing the MS data of HFGF-CD59 expressed in the HEK293-*HFGF-CD59* cells. Specifically, we analyzed precursor ions of *m/z* 912.32^3+^ (MW 2733.96, corresponding to HFGF-CD59 C-terminal peptide plus GPI containing three Man, two EthN-P and HexNAc). Indeed, in addition to fragment ions of *m/z* 1152^+^, and 1948^+^, other fragment ions *m/z* 528^+^ corresponding to P-Man-(EthN-P-) Man were found in some of the precursor ions of *m/z* 912.32^3+^ (Figure 5B), indicating that the C-terminal peptide of CD59 is attached to Man2-EthN-P, with the structure, peptide-EthN-P-(Man-)Man-(EthN-P-)(HexNAc-)Man-GlcN-PI (Figure 5A). These fragment ions were present in 2 out of 20 of precursor ions with MW 2733.0, which have three Man and two EthN-P (Table S4).

**Figure 5.**
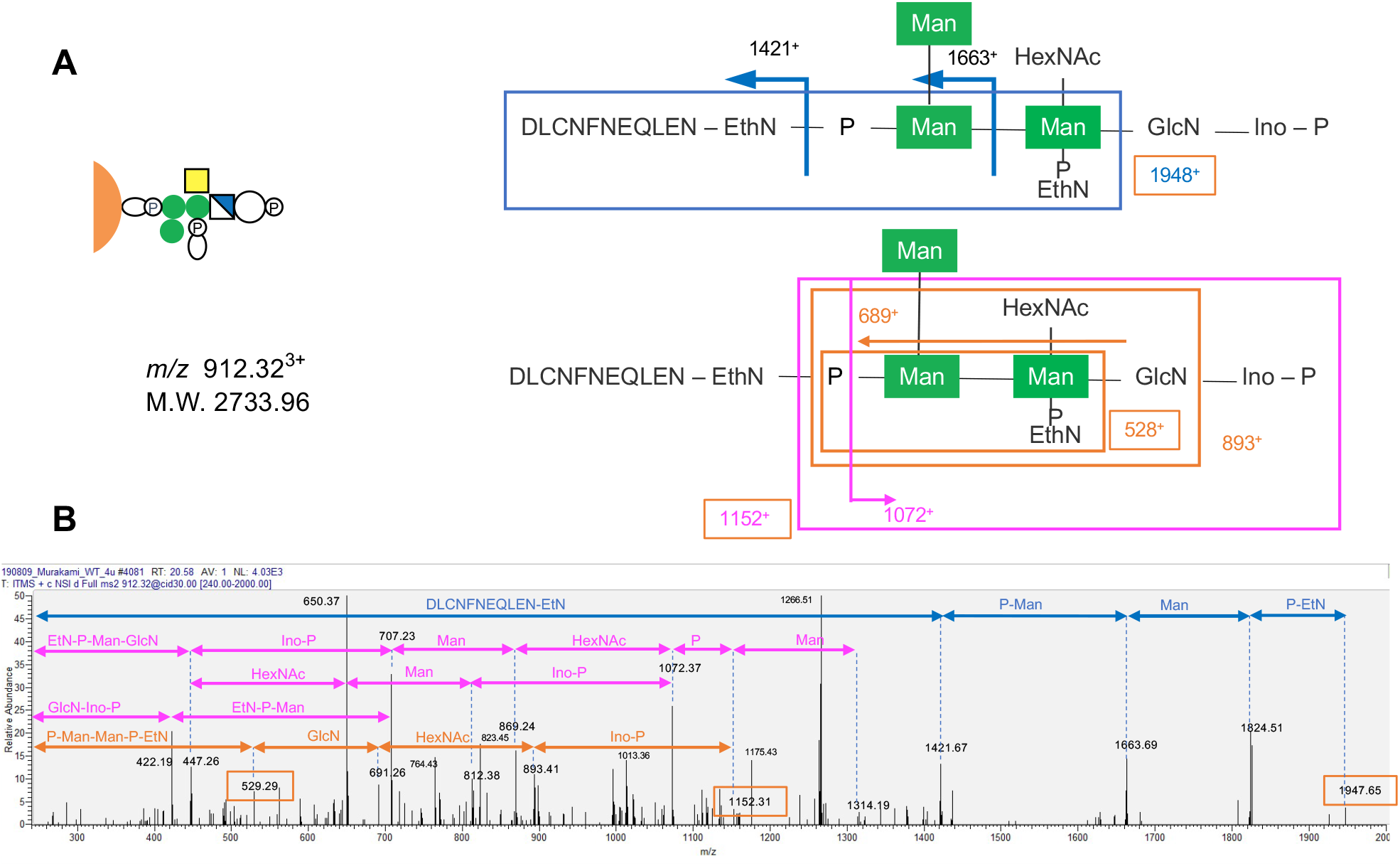
Protein attachment to EthN-P linked to the second mannose found in a small fraction of CD59 expressed in wild type HEK293 cells. A. Schematic MS/MS profile of CD59 C-terminal eleven amino acids linked to the second mannose of a GPI glycan. M.W. was calculated from the value of *m/z*. B. MS/MS analysis of a precursor ion of *m/z* 912.32^3+^. Fragment ions of *m/z* 529^+^,689^+^, 893^+^, 1152^+^ and 1948^+^ (diagnostic for protein attachment to EthN-P on the second mannose) were found in approximately 10% of precursor ions of *m/z* 912^3+^ and1367.5^2+^ (11 amino acids plus GPI consisting of three mannoses, two EthN-P, GlcN, HexNAc, inositol and phosphate), indicating protein attachment to EthN-P on the second mannose.

### Expression of several GPI-APs are PIGG dependent and preferentially bridged through Man2-linked EthN-P

The results showing that approximately 10% of HFGF-CD59 are bridged through Man2-linked EthN-P in HEK293 cells transfected with the tagged *HFGF-CD59* gene (Figure 5 and Table S4), also suggest the possibility that some GPI-APs are preferentially linked in this way (Figure S4). If this is true, the expression of these proteins might be lower in *PIGG*-KO cells than in wild type cells. To test this hypothesis, GPI-APs were isolated from wild type and two clones of *PIGG*-KO HEK293 cells, by a combination of repeated Triton X114 phase separation and a PI-PLC treatment (Figure 6).

**Figure 6.**
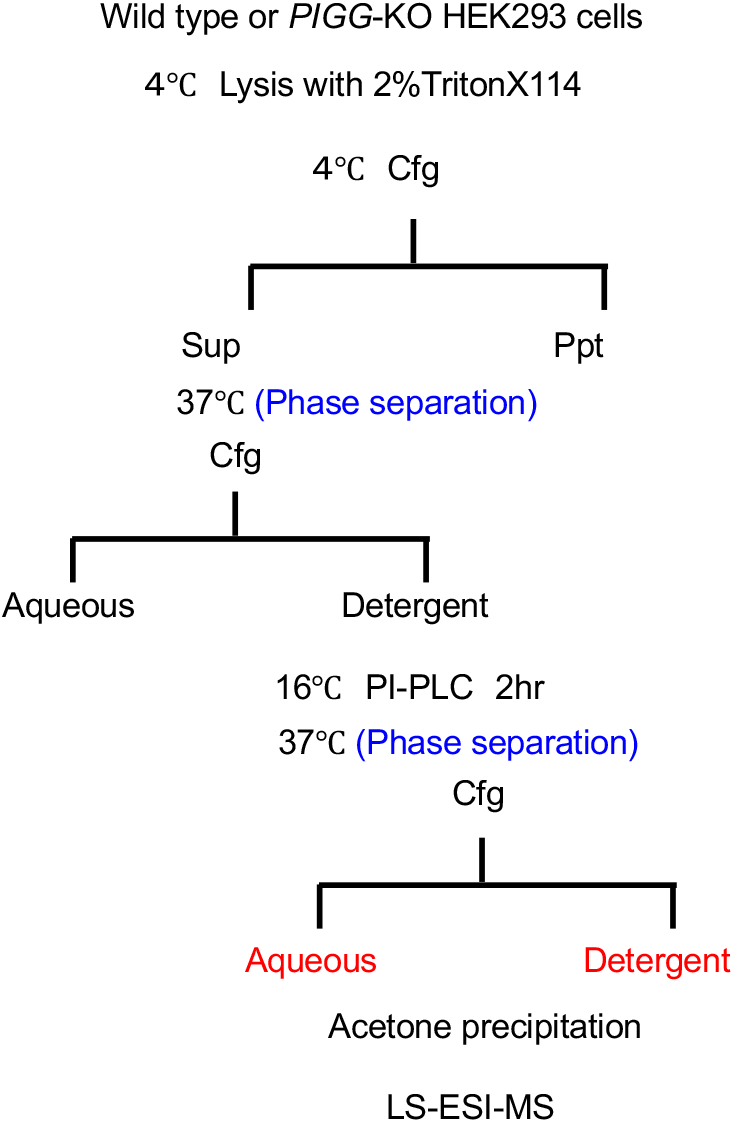
Flowchart of method for purification and fragmentation of GPI-APs for MS analysis. Cells are lysed at 4°C in Triton X114, followed by low-speed centrifugation and warming up of supernatant to 37 °C for phase separation. The detergent phase is collected and cold buffer (10 mM Tris-HCl, pH 7.4, 150 mM NaCl, 5 mM EDTA, proteinase inhibitor) added to dilute the TritonX-114, followed by PI-PLC treatment at 4°C for 90 minutes, and a second phase separation. PI-PLC-sensitive GPI-APs fragments stay solubilized in the aqueous phase, while PI-PLC-resistant GPI-APs stay into the detergent phase. GPI-APs fragments are concentrated by acetone precipitation and centrifugation and subjected to MS analysis.

The isolated proteins were then analyzed by LC-ESI-MS/MS and a total of 36 GPI-APs identified (Table S1). Of these, seven GPI-APs (tissue-nonspecific alkaline phosphatase (ALPL), ciliary neurotrophic factor receptor (CNTFR), contactin1 (CNTN1), melanotransferrin (MELTF), ecto 5’-nucleotidase (NT5E or CD73), reversion-inducing-cysteine-rich protein with kazal motifs (RECK), and reticulon 4 receptor like 2 (RTN4RL2)), were lower in the two *PIGG*-KO cloned cells than in wild type HEK293 cells (Table S1), as determined by comparative measurements using the Scaffold4 software (Searle, 2010).

To confirm these results, cell surface levels of these GPI-APs were also analyzed by FACS. Observed levels of NT5E (43.8±11.0% of wild type level, n=4) and EphrinA5 (53.7±10.0%, n=3) were lower in the *PIGG*-KO (Figure 7A). No changes in the levels of CD109, DAF or CD59 were observed (Figure 7A). The reduced level of NT5E, but not that of EphrinA5, on *PIGG*-KO cells was rescued by transfection of *PIGG* cDNA (Figure 7B). Furthermore, the cell surface levels of CD109, DAF, CD59, MELTF and prion protein (PRNP) on *PIGG*-KO cells did not increase upon transfection of *PIGG* cDNA (Figure 7B). These results show that the GPI-APs, which were rescued by transfection of *PIGG* cDNA, are PIGG dependent and preferentially bridged through the Man2-linked EthN-P.

**Figure 7.**
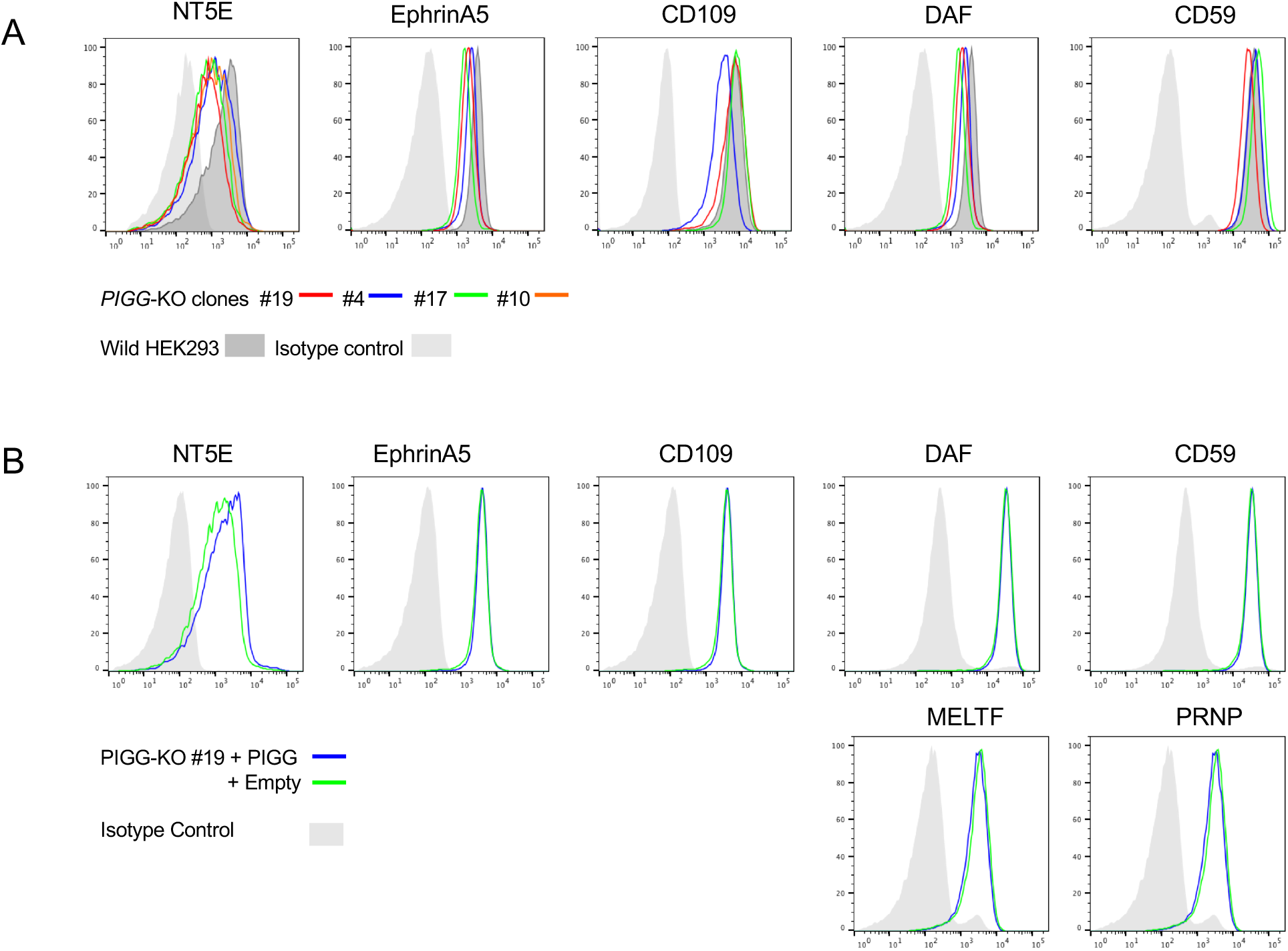
Expression levels of various GPI-APs in *PIGG*-KO cells compared to levels in the wild type HEK293 cells, determined by FACS analysis. A. Expression levels of various GPI-APs in different clones of *PIGG*-KO HEK293 cells. Fluorescence intensities of NT5E in *PIGG*-KO clones #4, #10, #17, #19 and wild type cells are 575, 910, 696, 530 and 1548 while those of EphrinA5 in clones #4, #17, #19 and wild type cells are 1431, 2080, 1868 and 3336, respectively. B. Expression levels of GPI-APs in *PIGG*-KO clone #19 cells stably transfected with *PIGG* cDNA compared to levels in empty vector transfected cells. Fluorescence intensities of NT5E in *PIGG* cDNA and in vector transfected cells are 1332 and 886, respectively.

To further test if expression of other GPI-APs are similarly PIGG-dependent, various GPI-APs genes were HA-tagged and cloned using the *pME* expression vector. Genes of fifty GPI-APs, known to be expressed in the brain, were selected and cloned for this work, and each recombinant plasmid co-transfected with *PIGG* cDNA, or an empty vector as control, into *PIGG*-KO HEK293 cells, followed by FACS analysis of the proteins expressed on the cell surface (Figure S5A and S5B). Expression levels of 18 of the GPI-APs including ALP, CNTFR, CNTN1, RECK, NT5E, and glypican 3 (GPC3), were clearly lower in the *PIGG-*KO cells as compared to the *PIGG* cDNA rescued transfectants (Figure S5A), whereas those of 32 other GPI-APs were similar, or only slightly lower (Figure S5B). Repeated measurements of 14 out of the former group of 18 GPI-APs resulted in levels of expression between 28.3±7.6 % and 62.5±13.8 % in *PIGG-KO* cells compared to those in *PIGG* cDNA rescued cells, while levels of 4 GPI-APs in the latter group of 32 were between 68.0±6.0 % and 88.1±3.9 % (mean±SD) (Figure 8A). These results indicate that the expression of the 18 GPI-APs tested is partially PIGG dependent and the extent of PIGG dependency varies among the proteins. Of those, Netrin G2 (NTNG2) was the most PIGG-dependent, showing approximately 70% reduction in *PIGG*-KO cells (Figure 8B). In other words, this *PIGG* dependency indicates that many GPI-APs, to varying degrees, are preferentially linked to Man2-linked EthN-P.

**Figure 8.**
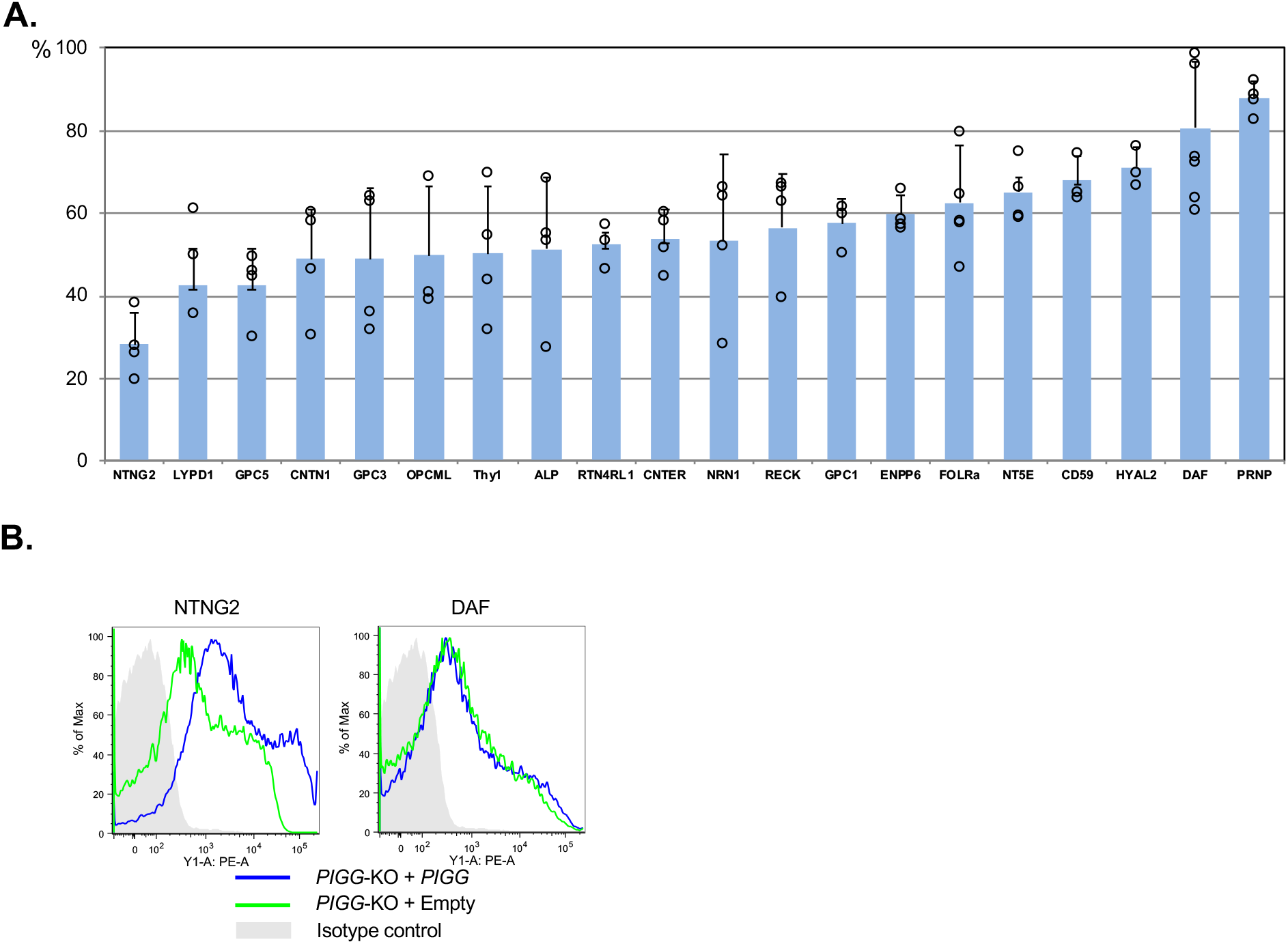
PIGG dependent expression of GPI-APs (FACS analysis). A. Expression levels of different GPI-APs in *PIGG*-KO HEK293 cells transiently co-transfected with *PIGG* cDNA or empty vector. Vertical bars represent % of levels of GPI-APs expression in non-rescued *PIGG*-KO cells relative to those in *PIGG* cDNA rescued *PIGG*-KO cells. B. FACS analysis of HA tagged *NTNG2* and FLAG tagged *CD55* co-transfected with *PIGG* cDNA or empty vector to *PIGG*-KO cells. Fluorescence intensity levels of NTNG2 in *PIGG*-KO cells and in *PIGG* rescued cells are 639 and 3426, while those of DAF in *PIGG*-KO cells and in *PIGG* rescued cells are 562 and 599, respectively.

To demonstrate attachment of these proteins to Man2-linked EthN-P by LC-MS/MS, we chose NT5E for analysis. HA tagged NT5E was affinity purified from the supernatant of PI-PLC treated Expi 293F cells transfected with N-terminally HA tagged NT5E, followed by SDS-PAGE and in-gel trypsin digestion (Figure S6). By LC-MS analysis, we found precursor ions corresponding to the C-terminal 2 amino acid (FS) peptide linked to GPIs having three Man and a HexNAc side chain (*m/z* 796.24^2+^; MW 1590.48) as well as some bearing Hex extension (*m/z* 877.26^2+^; MW 1752.52) (Figure 9 and Table S4). MS/MS analysis indicated not only the GPI diagnostic fragments, such as those with *m/z* 422^+^, 447^+^, 707^+^, 812^+^, 869^+^, 1072^+^ and 1314^+^, but also the fragments with *m/z* 484^+^, 949^+^, and 1152^+^. The fragment with *m/z* 484^+^ corresponds to P-Man-Man-P and those with *m/z* 949^+^, and 1152^+^ are bigger fragments containing P-Man-Man-P (Figure 9). They are specific to protein attachment to Man2-linked EthN-P. In addition, peptide-containing fragments (m/z 1227^+^and 762^+^) were detected, further confirming its structure. These fragment ions were present in all precursor ions with MW 1590 and 1752 (Table S4). These results indicated that NT5E expressed in wild type HEK293 cells has the preference to PIGG dependent structure.

**Figure 9.**
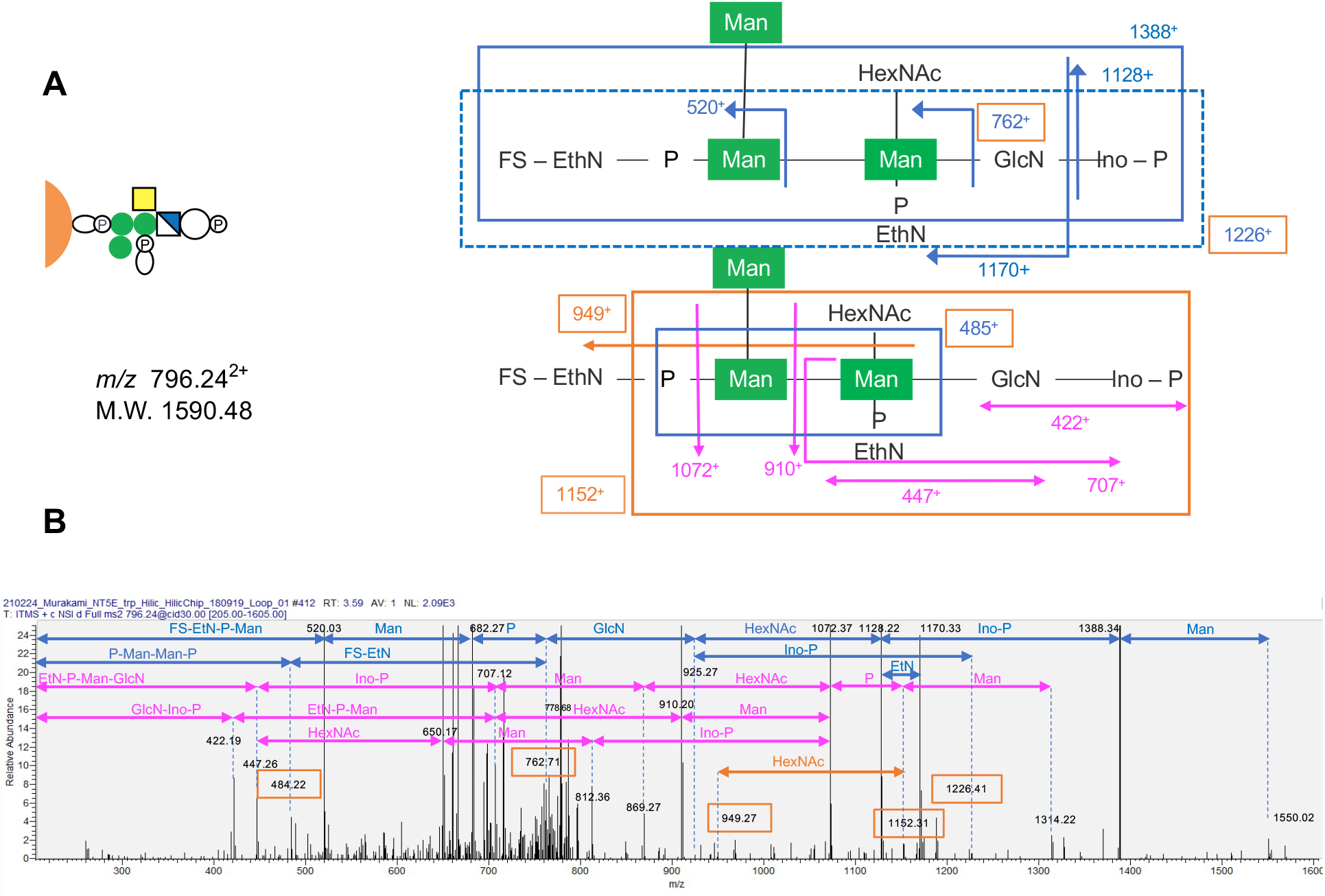
Protein attachment to EthN-P linked to the second mannose found in the majority of NT5E expressed in wild type HEK293 cells. A. Schematic MS/MS profile of NT5E C-terminal two amino acids (FS) linked to EtN-P on the second mannose of a GPI glycan. M.W. was calculated from the value of *m/z*. B. MS/MS analysis of a precursor ion of *m/z* 796.24^2+^ (two amino acids plus GPI consisting of three mannoses, two EthN-P, GlcN, HexNAc, inositol and phosphate). Fragment ions of *m/z* 485^+^, 762^+^, 949^+^, 1152^+^, and 1227^+^(diagnostic for protein attachment to EthN-P on the second mannose) were found in all precursor ions of M.W. 1590, indicating protein attachment to EthN-P on the second mannose is the major form in NT5E.

### Establishment of a more sensitive and simpler functional assay for PIGG mutations in individuals with IGD

The protocol shown above for measuring PIGG dependent residual expression of some GPI-APs (Figure 1C), lead us to propose a method without using radioactive material, for assaying *PIGG* mutations. Previously, the activity of PIGG could only be determined by metabolic labelling of *PIGG* cDNA-transfected cells with radioactive mannose (^14^C-mannose) and measuring the GPI intermediate bearing EthN-P on Man2 by thin layer chromatography and phosphorimaging (Makrythanasis et al., 2016). This method is time consuming and requires special facilities and training in radioisotope use. An example of the techniques we used are the ones for detecting residual expression of GPI-APs in *PIGO*-KO cells. PIGG activity on GPI-APs can be measured by transfecting wild type or mutant *PIGG* cDNA into *PIGO/PIGG-*DKO cells, followed by FACS analysis (Figure S6). These are standard cellular and biochemical techniques, being relatively simple and sensitive, and may be further adapted for the prognosis of individuals with *PIGG* mutations.

## Discussion

The core structure of GPI-APs is conserved among eukaryotes and it has been postulated that these proteins are always attached via EthN-P on Man3. In this study, we demonstrate the presence of a structural feature in GPI-APs not described before, in which EthN-P on Man2 bridges GPI anchor to proteins. We initially found that the residual expression of GPI-APs in *PIGB*-KO cells and *PIGO*-KO HEK293 cells are PIGG dependent (Figures 1 and 2). GPIs in *PIGB*-KO cells, which have only two mannoses, are attached to proteins via EthN-P linked to Man2 (Figure 4). Furthermore, we found that GPI with three mannoses and attached to a protein via EthN-P on Man2, is normally present in wild type HEK293 cells (Figure 5).

It is not yet known whether this alternative mode of protein attachment to GPI occurs in non-mammalian organisms. In organisms such as *Trypanosoma brucei*, in which only the Man3 is modified by EthN-P (Ferguson et al., 1988), this alternative mode of attachment may not occur. It has been reported that GPI core of *Entamoeba histolytica* proteophosphoglycan contains two mannoses and one galactose (Moody-Haupt et al., 2000). The authors suggest that the galactose might act in place of the Man3 and the bridging EthN-P might be linked to the galactose. In view of our findings, an alternative interpretation is that the galactose is only a side chain and the bridging is through EthN-P linked to Man2. Similarly, various GPIs purified from *Babesia divergens*, a protozoan parasite, contain a maximum of two mannoses (44). Further analyses of the GPI structure of GPI-APs on these protozoan parasites are warranted.

We used HFGF tagged CD59 to demonstrate protein attachment to EthN-P on Man2. Approximately 10% of the tagged CD59 overexpressed in wild type HEK293 cells had the alternative mode of attachment to GPI, as estimated by MS data analysis. Using a newly developed method of GPI-AP isolation combined with a proteomic analysis (Figure 6), we detected 36 GPI-APs in HEK293 cells (Table S1). Seven of them, ALPL, CNTFR, RECK, CNTN1, MELTF, NT5E/CD73, and RTN4RL2, were clearly decreased in two independent clones of *PIGG*-KO cells (Table S1). Decreased expression of NT5E in *PIGG*-KO cells was rescued by *PIGG* cDNA transfection (Figure 7). Among 50 GPI-APs known to be expressed in brain cells, many were highly, albeit not completely, PIGG-dependent (Figure 8), indicating that the alternative mode of GPI-anchor, bridged by Man2-linked EthN-P, is not a rare exception but occurs widely in many GPI-APs and to various extent.

The present results suggest causal relationship of GPI alternative bridging to *PIGG* mutations in IGD. Bi-allelic loss of function in *PIGG* mutations, found in individuals with IGD, may cause null or greatly decreased function of PIGG (Makrythanasis et al., 2016). It is likely therefore, that levels of a number of GPI-APs on the affected individuals’ cells are decreased. Consistent with this, lower levels of several GPI-APs in fibroblasts from two siblings with a homozygous nonsense *PIGG* variant, have been reported (Tremblay-Laganière et al., 2021; Zhao et al., 2017). Reduced levels of specific GPI-APs may result in the clinical symptoms associated with *PIGG*-IGD, such as developmental delay, intellectual disability, seizures and hypotonia. Among GPI-APs potentially decreased in PIGG-defective neuronal cells, is NTNG2, which is highly dependent upon PIGG (Figure 8). Recently, loss-of-function mutations in NTNG2, which is selectively expressed in presynaptic region, were found in several families, whose homozygous members showed global developmental delay, severe intellectual disability, muscle weakness, and behavioral abnormalities like Rett syndrome (Dias et al., 2019; Heimer et al., 2020). All these symptoms overlap with those of *PIGG*-IGD (Makrythanasis et al., 2016; Tremblay-Laganière et al., 2021; Zhao et al., 2017). Likewise, NT5E/CD73, which is widely expressed in brain tissue cells, might account for some of brain related abnormalities. This enzyme is known to convert AMP to adenosine. Functions of NT5E/CD73 in the central nervous system include roles in locomotion and behavior, memory and plasticity, sleep regulation, and thermoregulation (Minor et al., 2019). We successfully analyzed structures of GPI in NT5E expressed in wild type HEK293 (Epi293F) cells by mass spectrometry and showed that almost all were attached to EtN-P linked to Man2, the PIGG dependent structure (Figure 9 and Table S4). This is consistent to the fact that the PIGG deficient patients’ fibroblasts showed the decreased expression of NT5E (Tremblay-Laganière et al., 2021; Zhao et al., 2017). In *PIGG* KO cells, NT5E/CD73 expression was only slightly decreased, suggesting that EtN-P on the third Man can work as an alternative bridge in *PIGG* KO cells. We also tried to determine GPI structures in NTNG2 but unfortunately, repeated mass spectrometric analysis was unsuccessful and did not obtain the structural information.

GPI transamidase complex, consisting of PIGK, GPAA1, PIGT, PIGU, and PIGS (Eisenhaber et al., 2014; Hong et al., 2003; Ohishi et al., 2001; Yu et al., 1997), catalyzes the cleavage of the C-terminal signal peptide of GPI precursor proteins between the ω and ω+1 amino acids, and the formation of an amide bond between the ω amino acid carboxyl group and EthN of GPI. The catalytic subunit, PIGK is a cysteine protease that cleaves the C-terminal peptides and forms a carbonyl intermediate (Yu et al., 1997). GPAA1 has sequence homology to an M28 family peptide-forming enzyme, which catalyzes the formation of an amide bond between the carboxyl group of ω amino acid and amino group of EthN (Eisenhaber et al., 2014).

Other crucial questions on how this GPI transamidase complex functions still remain. For instance, in wild type cells, how it targets EthN-P to Man2 or Man3? It might be that some precursor proteins are conformationally more fit to the Man2-linked EthN-P. Alternatively, it is also possible that the conformation of the enzyme complex changes depending upon the precursor protein, setting the target preference.

Because PIGG deficiency mainly causes neurological defects, it is possible that GPI-APs in brain have the alternative mode of GPI attachment happening more frequently than those in other tissues/organs. The question, what could be the functional difference between the two structures of brain GPI-APs, remains, which leads to other questions as to why and how PIGG deficiency causes the various neurological symptoms (Makrythanasis et al., 2016; Tremblay-Laganière et al., 2021; Zhao et al., 2017). Furthermore, there must be a reason why some proteins are expressed with this alternative structure. Each GPI-AP works as a receptor, a ligand, an adhesion molecule, or as an enzyme. It is very possible that binding partners end up with different binding affinities to the GPI-APs with the alternate structure. We found decreased levels of many GPI-APs in *PIGG*-KO cells. The decreased levels of these proteins in brain of PIGG deficient individuals, certainly result in altered biological processes and hence, causing their various symptoms.

In support of altered biological processes caused by PIGG deficiency, in yeast, *PIGG*-KO cells show growth defect and it is reported that the rate of ceramide replacement in the lipid moiety of GPI anchors is reduced, the transport of GPI-APs to the Golgi is delayed, and cell separation is disturbed (Fujita et al., 2004).

Soon after the attachment of GPI to a precursor protein, the acyl chain linked to the inositol ring is usually removed by the GPI-deacylase PGAP1 (Tanaka et al., 2004). If removal of the acyl chain is prevented, the GPI-AP becomes PI-PLC resistant. CD59 in *PIGO*-KO cells is completely resistant to PI-PLC, even with overexpression of *PGAP1* (results not shown), whereas in *PIGB*-KO cells it is partially sensitive, albeit less sensitive as compared to that in wild type cells. PI-PLC does not act on CD59 expressed in *PIGO*-KO cells most likely because PGAP1 does not remove acyl chain from inositol in GPI, due to Man2-linked EthN-P as protein attachment and Man3 lacking EthN-P. In contrast, CD59 on *PIGB*-KO cells is partially sensitive to PI-PLC. This suggests that even when GPI has Man2-linked EthN-P being used in protein attachment, if Man3 is absent, PGAP1 acts on inositol-linked acyl chain. It is possible that PGAP1 recognizes these parts of GPI around Man2 and Man3 during its deacylating action. From the proteomic analysis, it seems that PI-PLC sensitivity has no direct relation to the alternate structure of GPI-APs but rather to the inherent differences of PI-PLC sensitivity of different GPI-APs. It is also known that PI-PLC sensitivities of these proteins are different depending on their cell type origin. For instance, GPI-APs in red blood cells are completely resistant to PI-PLC as they are not deacylated from inositol by PGAP1 (Roberts et al., 1988). PGAP1 activity is reported to have close association with ER stress (Liu et al., 2018). Under ER stress conditions, misfolded GPI-APs accumulate in the ER and deplete the available PGAP1, resulting in the exit of normal GPI-APs from the ER without being processed by PGAP1 (Liu et al., 2018). What structure is recognized by PGAP1 and how PGAP1 activity is regulated under non-stressed conditions, as well as the resulting GPI-APs sensitivities to PI-PLC under these conditions, are worth being clarified. Lastly, with the techniques we used for obtaining data on PIGG related effects, we propose a method to assess PIGG activity that is simpler and more sensitive. This new method can be further developed for more easily and accurately determine pathogenic effects of *PIGG* gene variants found in individuals with IGD.

In conclusion, we demonstrated that some GPI-APs are anchored through the EthN-P on Man2 of GPI, instead of the postulated Man3, and suggest that this newly discovered alternative mode of GPI attachment may be especially relevant in neurological development and functions.

## Methods

### Knockout (KO) cells

*PIGO-, PIGB-*, and *PIGG*-KO cells were generated from HEK293 cells using the CRISPR/Cas9 system (Cong et al., 2013). *PIGO/PIGG* and *PIGB/PIGG* double knockout (DKO) cell clones were generated by knocking out *PIGG* in cloned *PIGO-* and *PIGB-*KO cells, respectively. *PIGB/PIGZ* -DKO cells were generated by knockout of *PIGZ* in the cloned *PIGB*-KO cells. A pX330 plasmid for expression of the human codon-optimized *Streptococcus pyogenes* (Sp) Cas9 and chimeric guide RNA was obtained from Addgene (Cambridge, MA). The seed sequences for the SpCas9 target site in target genes are shown in Table S2. A pair of annealed oligos designed according to these sequences are inserted into the BbsI site of pX330. HEK293 cells were transfected with pX330 containing the target site using Lipofectamine 2000 (Invitrogen, Carlsbad, CA). Knockout clones were obtained by limiting dilution and KO confirmed by sequencing the target sites in the genome DNA and by flow cytometric analysis of GPI-APs. For the rescue experiments, N or C-terminal tagged human *PIGO, PIGB, PIGG*, and *PIGZ* cDNAs were cloned into the mammalian expression vector, pMEpuro18sf+ (Kitamura et al., 1991), generating pME-puro-3HA-hPIGO, pME-puro-3HA-hPIGB, pME-puro-hPIGG, pME-puro-hPIGZ-3HA, respectively (Table S3).

### Flow cytometry

Surface expression levels of GPI-APs were determined by staining cells with mouse anti-decay accelerating factor (DAF; clone IA10), -CD59 (clone 5H8) or -HA (clone HA-7, Sigma-Aldrich) followed by a PE-conjugated anti-mouse IgG antibody (BD Biosciences, Franklin Lakes, NJ). Cells were examined in the flow cytometer MACSQuantVYB (Miltenybiotec, Germany) and data analyzed with the Flowjo software (v9.5.3, Tommy Digital, Tokyo, Japan).

### Generation of PIGB-KO cells expressing His-FLAG-GST-FLAG tagged CD59 (HFGF-CD59)

Wild type HEK293 cells were stably transfected with pME-puro-HFGF-CD59 and highly expressor cells were sorted after labeling with anti-CD59. *PIGB-KO* cell clones were generated from these cells using CRISPR/Cas9 system, as described above. From the HFGF-CD59-expressing *PIGB*-KO cells, *PIGB/PIGZ*-DKO cells were further generated.

### Purification of HFGF-CD59

Cells (1×10^8^) stably expressing HFGF-CD59 were treated with 0.05 units/ml of phosphatidylinositol-specific phospholipase C (PI-PLC) (ThermoFisher, P6466) in 10 ml of PI-PLC buffer (Opti-MEM containing 10mM HEPES-NaOH, pH 7.4, 1 mM EDTA and 0.1% BSA) at 37 °C for 1 h. After centrifugation at 400 g for 5 minutes, the supernatant was loaded to a column of 500 μl of Glutathione Sepharose 4B at 4 °C. After washing repeatedly with a total of 10 ml PBS, protein was eluted with 5 ml of elution-buffer A (PBS containing 30 mM HEPES-NaOH, pH7.4, and 20 mM reduced glutathione). Protein in the eluted solution was precipitated with four volumes of acetone at -80 °C for 1h. After centrifugation at 13,400 x g for 30 min at 4 °C, the pellet was air dried dissolved in NuPAGE LDS Sample Buffer with reducing agent (Invitrogen). The protein pellet was further purified by electrophoresis in the Bolt 4-12% Bis-Tris Plus gel (Invitrogen), the gel stained with Imperial Protein Stain (Thermo Fisher Scientific) and a gel slice with the corresponding HFGF-CD59 protein excised.

### Purification of HA-NT5E

10μg of HA-tagged NT5E expressing plasmids (pME HA-NT5E) were transiently transfected with ExpiFectamine into 2.5×10^7^ Expi293F cells in 10ml medium according to manufacturer’s protocol (Gibco Expi293 Expression System, Thermo Fisher). One day after transfection, 0.05 units/ml of PI-PLC was added to the medium and cells were further incubated with shaking for three days. After centrifugation at 400 x g for 5 min, followed by filtration with the 0.45*μ*m filter, the supernatant was incubated for o/n with 40*μ*l of anti-HA agarose beads (rat anti-HA, 3F10 Roche) at 4 °C. After washing three times with PBS, protein was eluted with 80*μ*l of sample buffer and applied to SDS-PAGE. The gel stained with Imperial Protein Stain and a gel slice with the corresponding HA-NT5E protein excised.

### Mass spectrometric (MS) analysis

The gel slice with the HFGF-CD59 protein was reduced with 10mM dithiothreitol (DTT), followed by alkylation with 55mM iodoacetamide, and in-gel digestion with trypsin. The resultant peptides were subjected to MS analysis by LC-ESI system. Data were obtained by nanocapillary reversed-phase LC-MS/MS using a C18 column (0.1 × 150 mm) on a nanoLC system (Advance, Michrom BioResources) coupled to an LTQ Orbitrap Velos mass spectrometer (Thermo Fisher Scientific). The mobile phase consisted of water containing 0.1% formic acid (solvent A) and acetonitrile (solvent B). Peptides were eluted with a gradient of 5–35% solvent B for 45 min at a flow rate of 500 nl/min. The mass scanning range of the instrument was set at *m/z* 350–1500. The ion spray voltage was set at 1.8 kV in the positive ion mode. The MS/MS spectra were acquired by automatic switching between MS and MS/MS modes (with collisional energy set to 35%). Helium gas was used as collision gas. In-house MASCOT Server (Matrix Science) and Xcalibur Software (Thermo Fisher) were used for analysis of mass data. In the MS/MS profiles, those that contained characteristic fragments derived from GPI anchors, such as fragment ions of *m/z* 422^+^ and 447^+^, were selected, and fragments in the selected profiles were assigned to determine the GPI structures. Based on the profiles of the MS/MS fragments, the peak areas of the parental MS fragments corresponding to predicted GPI-peptides were measured in total ion chromatogram and the ratio in the total GPI peptide fragments calculated.

The gel slice with the HA-NT5E protein was reduced with 10mM dithiothreitol (DTT), followed by alkylation with 55mM iodoacetamide, and in-gel digestion with trypsin. The resultant peptides were subjected to MS analysis by LC-ESI system using a hydrophilic column. The samples were purified with Nu-Tip polyHydroxyethylA (HILIC) tips (GlySci) according to the manufacturer’s protocol before measurements. Purified peptides were then dissolved in 0.1% formic acid for measurement. Data were collected in positive ion mode on a nanoLC system using an EX-Nano Inert Sustain Amide capillary (0.1 × 150 mm; particle size, 3 μm) coupled to the same mass spectrometer. Measurement samples were injected onto the column by loop injection method. The mobile phase was the same as above, and peptides were eluted by. The same MS parameters were used as described above. The obtained mass data were analyzed by Xcalibur (Proteome Discoverer1.4 v. 1.4.0.288), and peptides were identified by MASCOT v. 2.7(2.7.0) (Matrix Science) using the SwissProt_2021_03 database (565,254 sequences; 203,850,821 residues; taxonomy: *Homo sapiens* (human) (20,387 sequences)). Protease specificity was set for trypsin (C-term, KR; Restrict, P; Independent, no; Semispecific, no; two missed and/or nonspecific cleavages permitted). Fixed modification considered was carbamidomethyl cysteine, and variable modifications considered were acetyl N terminus, N-terminal Gln to pyro-Glu, and oxidation of methionine. For quantification, the intensities of the top three peaks (monoisotopic peak + 0.5 or 0.33 + 1.0 or 0.66 Da for divalent or trivalent ions, respectively) corresponding to GPI-containing peptides in the MS spectra were calculated.

### *Proteomics analysis of* wild or *PIGG*-KO HEK293 cells

Wild type and two *PIGG*-KO clones (#10, #4) of HEK293 cells (5×10^6^ cells each) were lysed in 500 μl of lysis buffer (10 mM Tris-HCl, pH 7.4, 150 mM NaCl, 5 mM EDTA, protease inhibitor cocktail) containing 2% Triton X-114 (Nakalai Tesque, Kyoto, Japan) for 30 min on ice. After centrifugation at 15,000 rpm at 4°C for 15 min, the supernatant was incubated for 10 min at 37°C and aqueous and detergent phase separation was performed by centrifugation at 5,600g at 37°C for 7 min. After recovering the aqueous phase, 350 μl of lysis buffer added to the detergent phase and further incubated with PI-PLC (1 unit/ml) at 16 °C for 2h. Phase separation was performed and the aqueous phase combined with the previously recovered aqueous phase. Four volumes of acetone were added to the combined aqueous phase as well as to the detergent phase, and protein precipitation performed at -80 °C for 1h (Figure 6). Protein precipitate were pelleted by centrifugation at 13,400 x g for 30 min at 4 °C and the air dried. The air dried pellet was dissolved in 20 μl of 0.1% RapiGest (Waters) and reduced with 10 mM DTT, followed by alkylation with 55 mM iodoacetamide, digestion with trypsin and purification with a C18 tip (AMR, Tokyo, Japan). The purified peptides were subjected to nanocapillary reversed-phase LC-MS/MS analysis using a C18 column (10 cm x 75 um, 1.9 µm, Bruker Daltonics, Bremen, Germany) in a nanoLC system (Bruker Daltonics) connected to a timsTOF Pro mass spectrometer (Bruker Daltonics) and a modified nano-ESI source (CaptiveSpray; Bruker Daltonics). The mobile phase consisted of water containing 0.1% formic acid (solvent A) and acetonitrile containing 0.1% formic acid (solvent B). Linear gradient elution was carried out from 5% to 30% solvent B for 18 min at a flow rate of 500 nL/min. The ion spray voltage was set at 1.6 kV in the positive ion mode. Ions were collected in the trapped ion mobility spectrometry (TIMS) device over 100 ms and MS and MS/MS data collected over an *m/z* range of 100–1,700. During the collection of MS/MS data, the TIMS cycle was adjusted to 0.53 s and included 1 MS plus 4 parallel accumulation serial fragmentation (PASEF)-MS/MS scans, each containing on average 12 MS/MS spectra (>100 Hz) (Meier et al., 2015; Meier et al., 2018). Nitrogen gas was used as collision gas.

The resulting data was processed using DataAnalysis version 5.2 (Bruker Daltonics), and proteins identified using MASCOT version 2.6.2 (Matrix Science, London, UK) against the SwissProt database. Quantitative values were calculated with Scaffold4 (Proteome Software, Portland, OR, USA) for MS/MS-based proteomic studies (Searle, 2010).

### Determination of expression levels of GPI-APs in PIGG-KO cells

GPI-APs were HA-tagged by cloning their coding cDNA amplified from the human cDNA library or from Invitrogen plasmids, as described (Lee et al., 2020), into the mammalian expression vector (pME-puro) and co-transfected into the *PIGG*-KO cells with *PIGG* cDNA or the empty vector as control. The expression levels of the GPI-APs were determined by flow cytometric analysis

## Data availability

The mass spectrometry proteomics data have been deposited to the ProteomeXchange Consortium via the PRIDE (Vizcaíno et al., 2016) partner repository with the dataset identifier PXD021996 and PXD022032 (project name, “A novel structure of GPI anchored proteins”)

## Acknowledgments

We thank Junji Takeda (Osaka University) and Mike Ferguson (University of Dundee) for discussion, Luiz Shozo Ozaki (Virginia Commonwealth University) for discussion and manuscript editing, Shota Nakamura and Daisuke Motooka (Osaka University) for running the next generation sequencer, Hideki Nakanishi (Jiangnan University) for offering us the plasmids, Keiko Kinoshita, Saori Umeshita, and Kae Imanishi (Osaka University) for technical help. This work was supported by JSPS and MEXT KAKENHI grants (JP16H04753 and JP17H06422 for T. Kinoshita), a grant from Ministry of Health, Labor and Welfare and a grant from Practical Research Project for Rare/Intractable Diseases from the Japan Agency for Medical Research and Development (AMED) (21ek0109418h0003 for Y. Murakami).

## Author contributions

MI, PC, TK and YoM designed the study, MI, YuM, AN, YT and YoM acquired the data and conducted experiments. MI, PC, TK and YoM wrote the paper.

