## Supplemental materials for "Ethanolamine-phosphate on the second mannose is the preferential bridge for some of the brain GPI-anchored proteins"

**This PDF file includes:**

Figures S1 to S7
Tables S1 to S4
SI References

**Figure S1**

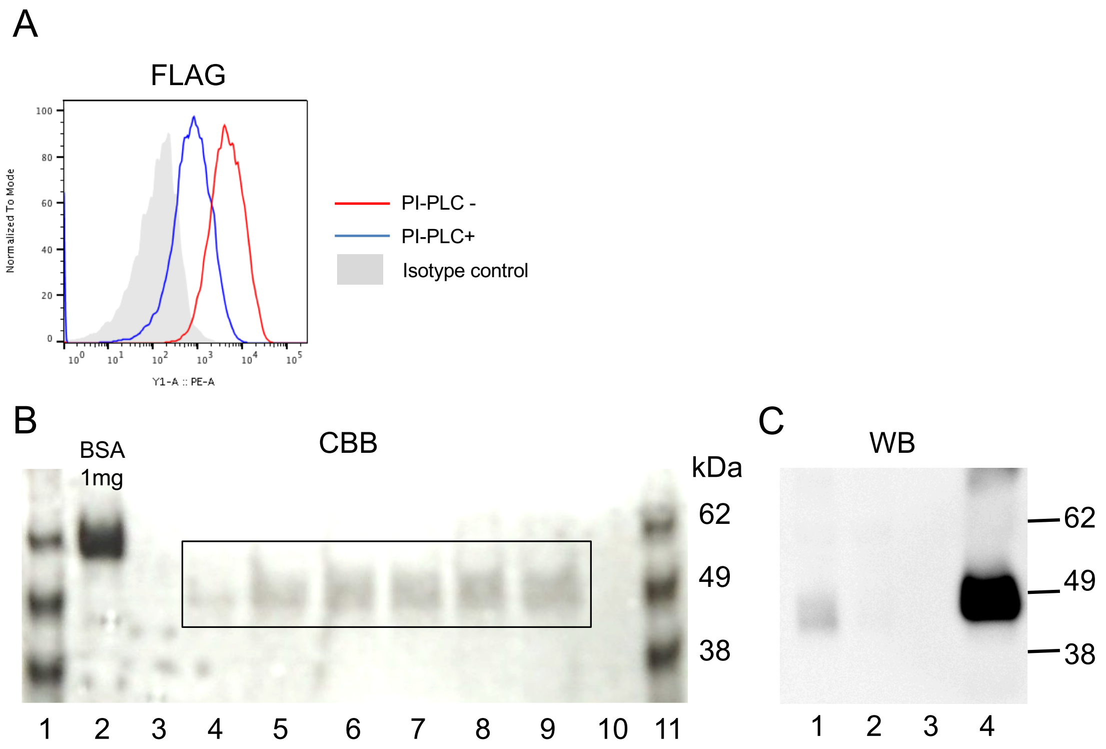


**Release of HFGF-CD59 synthesized in *PIGB*-KO HEK293 cells upon PI-PLC treatment.**

A. Analysis by flow cytometry of *PIGB*-KO HEK293 cells transfected with pME-puro-HFGF-*CD59* plasmid construct, with or without PI-PLC treatment. An estimated 80% of the *PIGB*-KO cells have their surface HFGF-CD59 released with PI-PLC treatment. B. SDS-PAGE analysis of glutathione affinity purified HFGF-CD59 expressed in *PIGB*-KO HEK293 cells and released by PI-PLC treatment (CBB, Coomassie Brilliant Blue). Lane 1 and 11, size markers (SeeBlue Plus2 Pre-Stained Protein Standard, Thermo Fisher Scientific); lane 2, BSA (bovine serum albumin, 1 mg); lane 4-9 (enclosed), HFGF-CD59. C. Western blot (WB) of the glutathione purified HFGF-CD59 using anti-GST antibody. Lane 1, aliquot of cell culture medium (6 μl of 11ml total) after PI-PLC treatment; lane 2, Unbound fraction (6 μl of 11ml total) after incubation with glutathione beads; lane 3, wash buffer solution (6 μl of 11ml total) of glutathione beads incubated with cell lysis extract; lane 4, elution buffer solution (5 μl of 120μl total) of glutathione beads incubated with cell lysis extract.

**Figure S2**

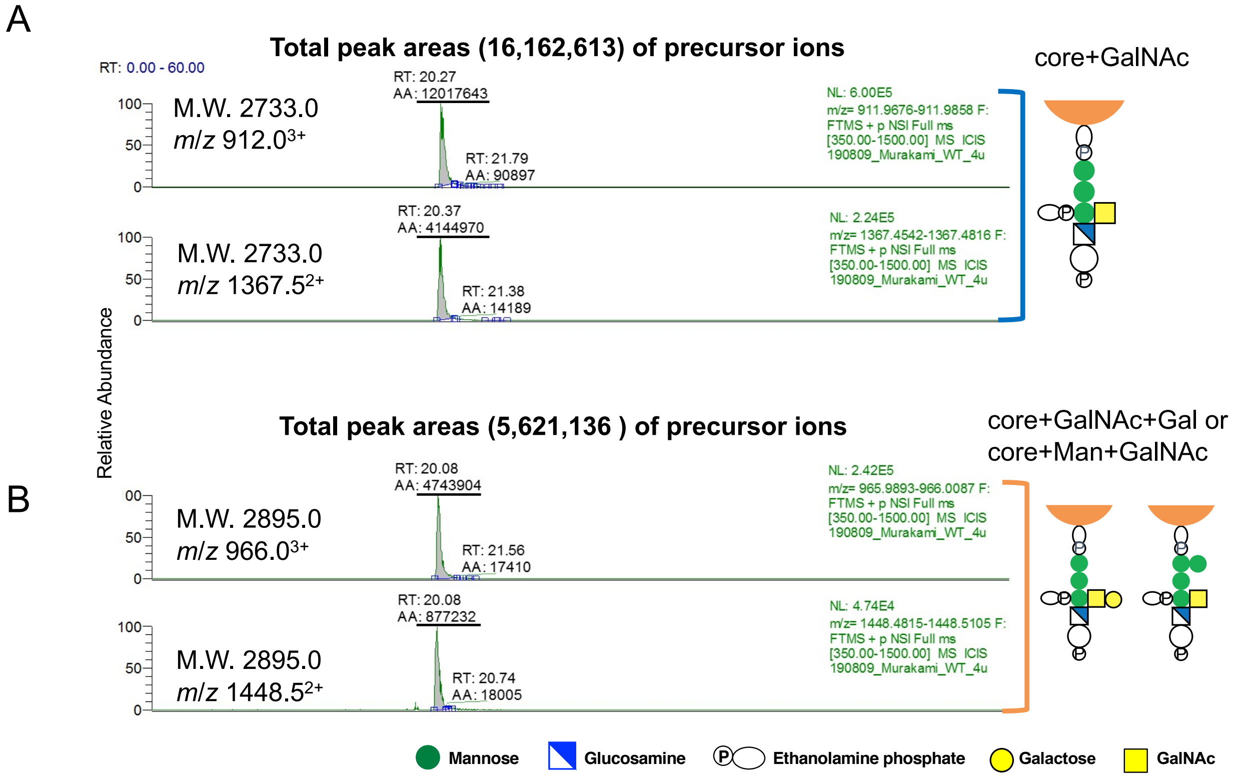


**Structures of GPI in HFGF-CD59 expressed in wild type HEK293 cells as determined by MS analysis.**

A. Total peak areas (16,162,613) of precursor ions *m/z* 912.0$^{3+}$ and 1367.5$^{2+}$ (bracket in blue), corresponding to a C-terminal peptide linked to GPI having the three-mannose core with GalNAc side chain (core + GalNAc). B. Total peak areas (5,621,136) of precursor ions *m/z* 966.0$^{3+}$ and 1448.5$^{2+}$ (bracket in orange) corresponding to GPI with the GalNAc side chain having either Gal extension or Man4. No precursor ions corresponding to any other GPI species were detected.

**Figure S3**

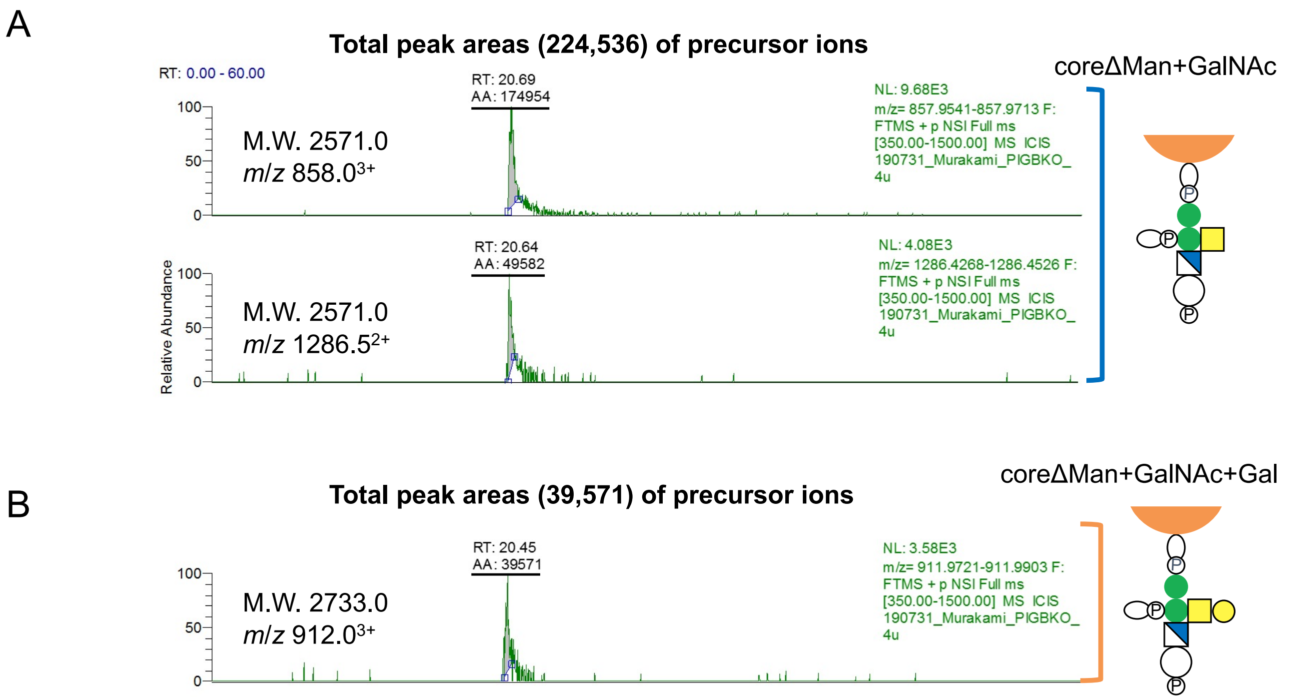


**Structures of GPI in HFGF-CD59 expressed in *PIGB*-KO HEK293 cells as determined by MS analysis.**

A. Total peak areas (224,536) of precursor ions *m/z* $858.0^{3+}$ and $1286.5^{2+}$ (bracket in blue), corresponding to a C-terminal peptide linked to GPI having the two-mannose core with GalNAc side chain (coreΔMan+GalNAc). B. Total peak areas (39,571) of precursor ion *m/z* $912.0^{3+}$ (bracket in orange) corresponding to GPI with the GalNAc side chain having Gal extension (coreΔMan+GalNAc+Gal).

### Figure S4

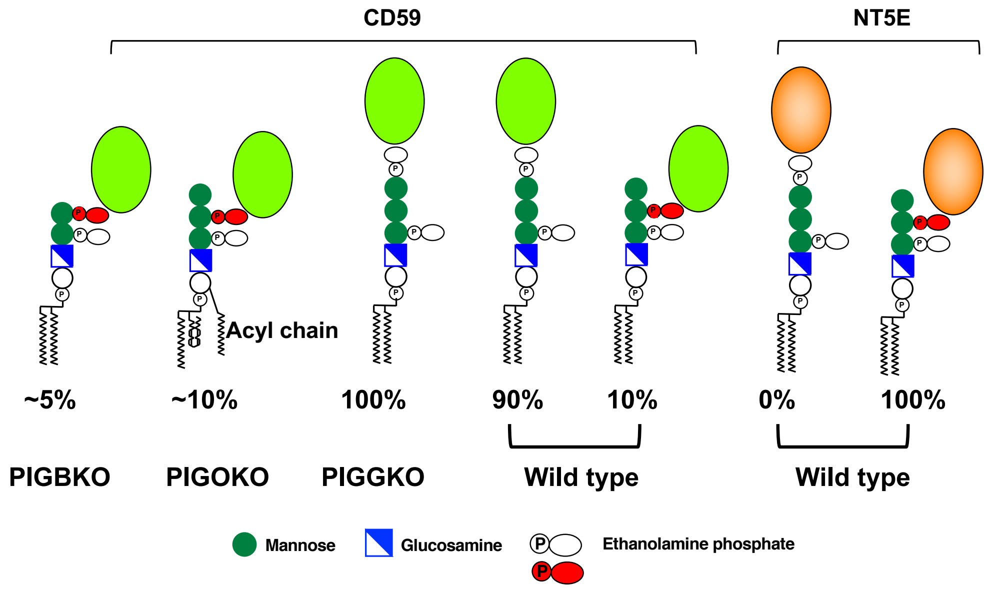


**Structure of the GPI moieties in CD59 and NT5E GPI-APs expressed in *PIGB-, PIGO-, PIGG*-KO (CD59) and wild type HEK293 (CD59 and NT5E) cells.** Proportion of CD59 with the GPI-Man2 in wild type HEK293 cells was calculated by MS analysis. Expression of CD59 in *PIGB-* and *PIGO*-KO cells was estimated by flow cytometry compared to wild type cells. Note: In *PIGG*-KO cells, CD59 with the GPI-Man2 structure is absent (100% GPI-Man3-CD59), which might happen with other GPI-APs, resulting in the neurological abnormalities, in people with *PIGG* mutations. As for NT5E, one of the *PIGG* dependent protein tested, the expression would be decreased compared to wild type cells in *PIGG* KO cells.

**Figure S5**

**A.**

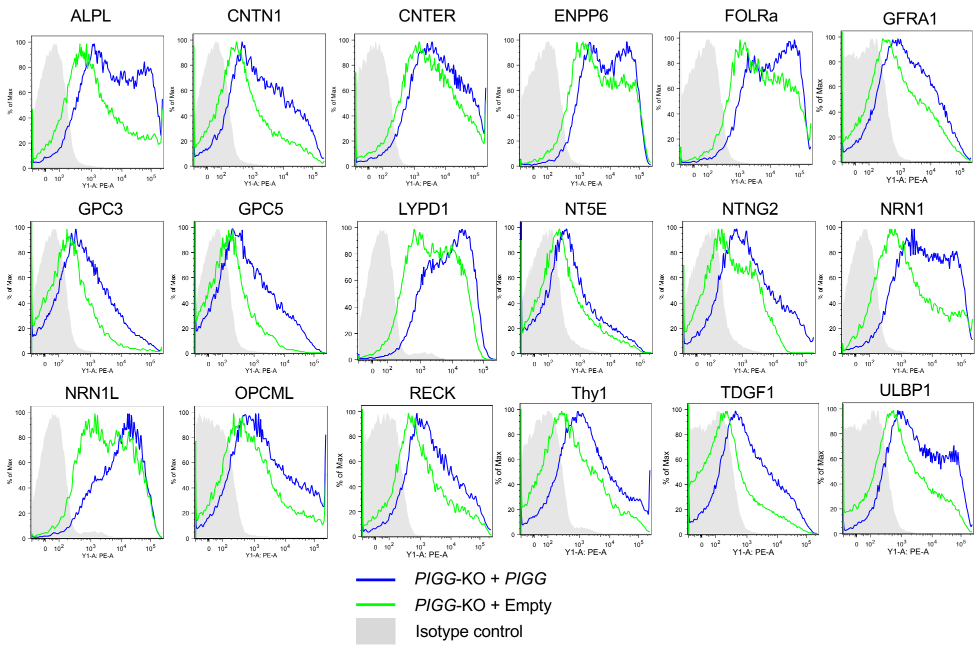


**B.**

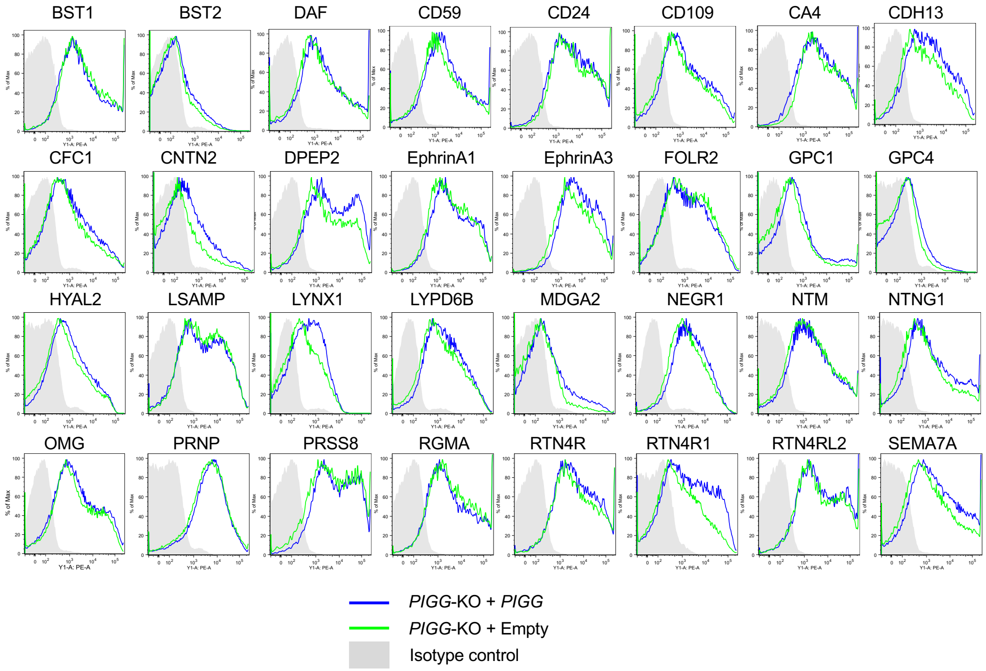


### PIGG dependent differential expression of GPI-APs

A. GPI-APs expression partially PIGG dependent. B. GPI-APs expression *PIGG* independent. Transient expression levels of each HA-tagged GPI-AP on HEK293 cells after co-transfection with *PIGG* cDNA or empty vector were assessed by flow cytometry after incubation with mouse anti-HA antibody and staining with PE conjugated anti-mouse IgG antibody.

**Figure S6**

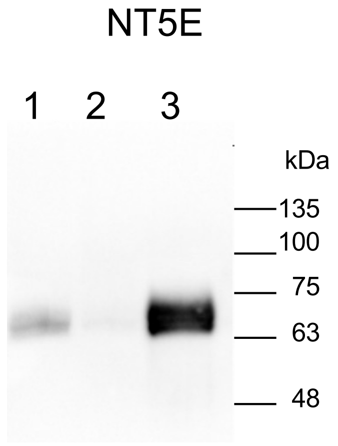


**Purification of HA tagged NTNG2 and NT5E proteins for Mass spectrometry**

HA-tagged NT5E expressing plasmids (pME HA-NT5E) were transiently transfected into Expi293F cells according to manufacturer's protocol (Gibco Expi293 Expression System, Thermo Fisher) and cells were incubated with PIPLC. Each protein was purified from the supernatant using HA-column (Roche). Lane1, supernatant; lane2, unbound; lane3, eluted protein.

**Figure S7**

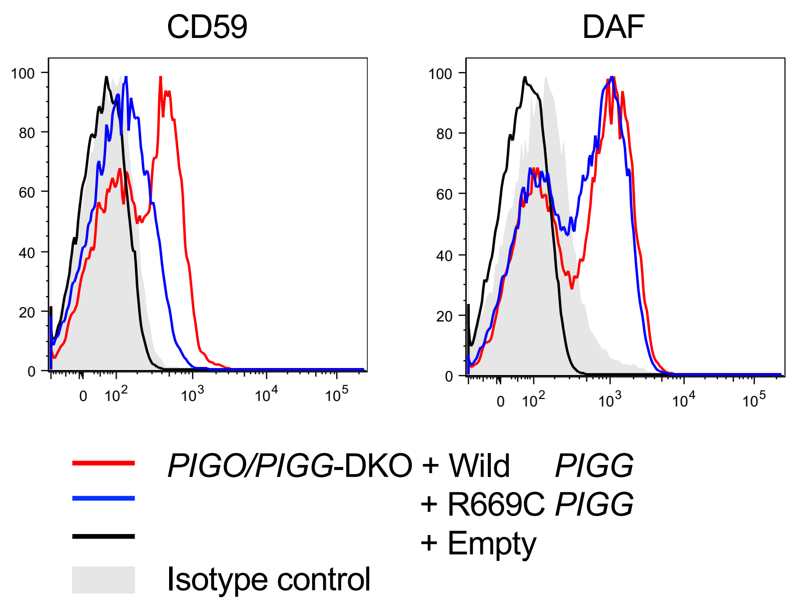


**Functional activity assay of R669C mutant *PIGG* cDNA on expression of GPI-APs in *PIGO/PIGG*-DKO cells (FACS analysis).**

*PIGO/PIGG*-DKO HEK293 cells transfected with wild type or R669C mutant *PIGG* cDNA with GST tagged C-terminal. The R669C-*PIGG* cDNA only partially restored the expression of CD59 and DAF (blue line) as compared with the restoration by the wild type *PIGG* cDNA (red line), suggesting that the activity of R669C-PIGG mutant was decreased.

### Table S1

Proteomics analysis of released proteins from PIPLC treated cells

| | Aqueous Phase | | Detergent Phase | | | | Aqueous Phase | | Detergent Phase | |
|---|---|---|---|---|---|---|---|---|---|---|
| Protein | WT | PIGGKO#4 | WT | PIGGKO#4 | | Protein | WT | PIGGKO#10 | WT | PIGGKO#10 |
| ALPL | 3.2048 | 0.79071 | 0 | 0 | | ALPL | 3.3218 | 0.97692 | 0 | 0 |
| BST2 | 4.0059 | 3.1628 | 5.193 | 9.0285 | | BST2 | 2.4914 | 0.97692 | 0 | 0 |
| CACNA2D1 | 12.018 | 15.814 | 16.877 | 18.057 | | CACNA2D1 | 6.6437 | 9.7692 | 4.6579 | 7.0913 |
| CD109 | 40.059 | 50.605 | 5.193 | 12.414 | | CD109 | 50.658 | 24.423 | 0 | 7.9777 |
| DAF | 22.433 | 25.303 | 0 | 0 | | DAF | 13.287 | 21.492 | 0 | 0 |
| CD58 | 3.2048 | 2.3721 | 1.2983 | 2.2571 | | | | | | |
| CD59 | 24.036 | 25.303 | 2.5965 | 3.3857 | | CD59 | 24.083 | 28.331 | 1.5526 | 2.6592 |
| CNTFR | 4.0059 | 0.79071 | 0 | 0 | | CNTFR | 2.4914 | 0 | 0 | 0 |
| CNTN1 | 29.644 | 18.186 | 1.2983 | 0 | | CNTN1 | 23.253 | 4.8846 | 0 | 0 |
| CPM | 4.8071 | 7.9071 | 0 | 2.2571 | | CPM | 5.8132 | 5.8615 | 0 | 0 |
| EFNA3 | 1.6024 | 1.5814 | 0 | 0 | | | | | | |
| EMC10 | 0.8012 | 1.5814 | 10.386 | 6.7714 | | | | | | |
| FOLR1 | 19.229 | 17.396 | 0 | 0 | | FOLR1 | 10.796 | 9.7692 | 0 | 0 |
| GAS1 | 6.4095 | 3.9535 | 1.2983 | 0 | | GAS1 | 4.9827 | 4.8846 | 0 | 0 |
| GFRA1 | 6.4095 | 8.6978 | 0 | 0 | | GFRA1 | 4.9827 | 0.97692 | 0 | 0 |
| GFRA2 | 4.0059 | 3.9535 | 0 | 0 | | GFRA2 | 4.9827 | 1.9538 | 0 | 0 |
| GPC1 | 2.4036 | 7.9071 | 0 | 0 | | GPC1 | 5.8132 | 7.8153 | 0 | 0 |
| | | | | | | GPC3 | 3.3218 | 0.97692 | 0 | 0 |
| GPC4 | 6.4095 | 10.279 | 0 | 0 | | GPC4 | 5.8132 | 4.8846 | 0 | 0 |
| HSPA5 | 56.083 | 52.187 | 5.193 | 3.3857 | | HSPA5 | 62.284 | 61.546 | 18.632 | 15.069 |
| HYAL2 | 2.4036 | 3.9535 | 0 | 0 | | HYAL2 | 2.4914 | 0.97692 | 0 | 0 |
| LYPD6B | 4.0059 | 2.3721 | 0 | 0 | | LYPD6B | 1.6609 | 1.9538 | 0 | 0 |
| MELTF | 40.861 | 26.884 | 0 | 0 | | MELTF | 39.031 | 18.561 | 0 | 0 |
| MICA | 1.6024 | 3.1628 | 0 | 0 | | | | | | |
| NCAM1 | 16.825 | 26.093 | 18.176 | 16.928 | | NCAM1 | 10.796 | 12.7 | 4.6579 | 14.183 |
| NT5E | 2.4036 | 1.5814 | 0 | 0 | | NT5E | 1.6609 | 0 | 0 | 0 |
| PGRMC1 | 4.0059 | 3.9535 | 25.965 | 32.728 | | PGRMC1 | 10.796 | 10.746 | 7.7632 | 24.819 |
| PLAUR | 8.0119 | 7.9071 | 0 | 0 | | PLAUR | 7.4741 | 6.8384 | 0 | 0 |
| PRNP | 8.8131 | 4.7442 | 2.5965 | 1.1286 | | | | | | |
| PRSS8 | 1.6024 | 0 | 0 | 0 | | | | | | |
| RECK | 5.6083 | 1.5814 | 0 | | | RECK | 3.3218 | 0 | 0 | 0 |
| RTN4R | 0.8012 | 1.5814 | 0 | 0 | | RTN4R | 4.1523 | 6.8384 | 0 | 0 |
| RTN4RL2 | 9.6143 | 7.1164 | 0 | 0 | | RTN4RL2 | 5.8132 | 1.9538 | 0 | 0 |
| SMPDL3B | 14.421 | 7.1164 | 0 | 0 | | SMPDL3B | 11.626 | 10.746 | 0 | 0 |
| ULBP2 | 2.4036 | 4.7442 | 0 | 0 | | | | | | |
| ULBP3 | 6.4095 | 3.9535 | 0 | 0 | | ULBP3 | 2.4914 | 4.8846 | 0 | 0 |

Quantitative value were calculated with Scaffold4 (Proteome Software, Portland, OR, USA) for MS/MS-based proteomic studies (1).

**Table S2**

Nucleotide sequences used for the CRISP/CAS9 guide RNAs

| Target gene | Sequence* |
|---|---|
| ***PIGB*** exon1 | 5’- GCAAGTGCGGAATGGAGCCGGGG - 3’ |
| ***PIGB*** exon5 | 5’- CAGAACCCTTACAAACACCATGG - 3’ |
| ***PIGG*** exon1 | 5’- GCGTAGCGATCGAGGTGCTAGGG - 3’ |
| ***PIGG*** exon2 | 5’- GCCCTACACAACTTACCTTGTGG - 3’ |
| ***PIGZ*** exon2 | 5’- GCACATAGCCCGTCTGCGGAAGG - 3’ |

*(Underlined nucleotides: PAM)

**Table S3**

Plasmid constructs used in this work

| | |
|---|---|
| *PIGO* cDNA | pME-puro-3HA-hPIGO |
| *PIGB* cDNA | pME-puro-3HA-hPIGB |
| *PIGZ* cDNA | pME-puro-hPIGZ-3HA |
| *PIGG* cDNA | pME-puro-hPIGG |
| *GST tagged PIGG* cDNA | pME-puro-hPIGG-GST |
| *HFGF-CD59* | pME-puro-HFGF (His-Flag-GST-Flag)-hCD59 |
| Empty vector | pME-puro-empty |

*HA, Human influenza hemagglutinin; h, human; GST, Glutathione S-transferase; puro, puromycin resistance gene

**Table S4**

Percentage of the precursor ions with the new structure (protein linked to the EtN-P on the second Man) among the GPI fragments derived from each GPI-AP

| Peptides | MW | GPI structure | GPI fragment(n) | New structure[*2] | Percent |
|---|---|---|---|---|---|
| Wild CD59 | 2733 | core[*1]+HexNAc | 20 | 2 | 10% |
| Wild NT5E | 1590 | core+HexNAc | 8 | 8 | 100% |
| | 1752 | core+HexNAc+Hex | 2 | 2 | 100% |

[*1] Core ; Peptide-EthN-P-Man-Man-(EthN-P-)Man-GlcN-Ino-P- or

Peptide-EthN-P-(Man)Man-(EthN-P-)Man-GlcN-Ino-P-

[*2] Precursor ions containing the fragments, which are specific to the new structure,

i.e., $485^+$, $528^+$, $762^+$, $949^+$, $1152^+$, $1226^+$
